# Tracking gene flow in the gut microbiome reveals phage activation and host-range expansion in response to diet

**DOI:** 10.64898/2026.10.01.756045

**Authors:** Saria McKeithen-Mead, Surya Tripathi, Theint Theint Thu, Steven Higginbottom, Adam Deutschbauer, Justin Sonnenburg, Kerwyn Casey Huang

## Abstract

Horizontal gene transfer (HGT) drives microbial evolution, but what determines when genes move between species within complex communities remains unclear. We developed a junction-resolved metagenomic framework using short-read sequencing to distinguish mobile genetic element activation from *de novo* transfer and track both longitudinally. In a defined human gut community colonizing mice, mobile element activation was widespread but successful transfer was rare. A prophage carried by invading *Phocaeicola vulgatus* transferred reproducibly to resident species when the community resisted invasion, but not when dietary perturbation allowed the invader to expand. Transfer required prophage activation and persistence of a permissive recipient and coincided with diversification of a phage receptor-binding protein. Dietary polysaccharides reproduced prophage-associated fitness costs *in vitro*, and rhamnogalacturonan I induced cell lysis consistent with productive prophage activation. The framework also detected interspecies transfer retrospectively in longitudinal human metagenomes. Thus, diet and community ecology can redirect gene flow through the gut microbiome.

## Introduction

Microbial communities regulate critical ecosystem processes, from biogeochemical cycling in soil and oceans to host-associated symbioses that support plant and animal life (*1-3*). In the mammalian gut, resident microbes influence host health by modulating immune function, metabolism, and pathogen resistance (*1*). Central to microbial adaptation in these dynamic environments is horizontal gene transfer (HGT), which enables the rapid acquisition of new functions without requiring *de novo* mutation (*4*). Mobile genetic elements (MGEs), including prophages, plasmids, and integrative conjugative elements, mediate most HGT (*5*), carrying both the machinery required for their own propagation and cargo genes that can reshape community composition and function (*6, 7*). Comparative genomics suggests that a substantial fraction of most bacterial genomes has been acquired horizontally (*8*), underscoring the profound evolutionary impact of HGT.

The mammalian gut is thought to be a hotspot for bacterial genetic exchange. High cell densities, close physical proximity, and the abundance of bacteriophages and conjugative elements create favorable conditions for transfer (*9*). At the same time, environmental perturbations such as dietary shifts (*10*), antibiotic exposure (*11*), and species invasion (*12*) continuously reshape community structure, potentially altering both opportunities for transfer and the selective pressures acting on transferred genes. Yet despite its recognized importance, the ecological conditions that promote HGT in complex communities remain poorly understood, largely because *de novo* transfer is difficult to track longitudinally in experimental systems that combine community complexity with the control needed to test ecological mechanisms.

Current methods for studying MGE dynamics, including Hi-C metagenomics (*13, 14*), long-read sequencing (*15*), single-cell sequencing (*16*), and cultivation-based approaches (*17-19*), can identify associations between MGEs and host genomes but generally provide static views of genetic exchange. These methods cannot readily distinguish active transfer from historical integration events or resolve the temporal sequence linking ecological perturbation, MGE activation, transfer, and selection. Experimental demonstrations of HGT *in situ* have often relied on antibiotic selection or engineered genetic markers, creating fitness landscapes that may not reflect native evolutionary pressures. As a result, the mechanisms connecting community ecology to HGT remain difficult to resolve.

Here, we developed a junction-resolved metagenomic framework that tracks MGE activation and transfer in real time from short-read sequencing data by distinguishing excised episomes from chromosomally integrated forms and identifying new integration events in recipient genomes. We applied this framework to a reproducible synthetic human gut community colonizing germ-free mice, using ecological invasion and dietary modulation to create contrasting community states. By combining longitudinal metagenomics with genome-resolved measurements of invader fitness, we connected MGE dynamics to their ecological and evolutionary consequences. Paradoxically, conditions that resisted invasion promoted prophage activation and horizontal transfer, whereas conditions that favored invasion suppressed both processes. We further found that specific dietary polysaccharides directly promote prophage activity, linking the dietary environment to prophage-mediated fitness costs and transfer. Prophage diversification during transfer revealed a dynamic sequence of molecular and ecological interactions accompanying gene transfer. Together, these findings establish a mechanistic pathway connecting diet, community structure, prophage activation, invader fitness, and horizontal transfer in the mammalian gut. Finally, application of our framework to a longitudinal human metagenomic dataset demonstrated that MGE transfer can also be tracked over time in naturally occurring human gut communities. More broadly, these results demonstrate how ecological context governs MGE dynamics in complex communities and establish junction-resolved metagenomics as a generalizable approach for tracking HGT across experimental and human microbial ecosystems.

## Results

### Diet-dependent community restructuring promotes microbial invasion

We established a defined 38-species community derived from the stool of a single human donor that reproducibly colonizes germ-free mice and remains stable under a standard diet (*20*) (SD, Methods) (**Fig. 1A,B**). To test how ecological invasion shapes horizontal gene transfer, we introduced a *Phocaeicola vulgatus* strain carrying a barcoded transposon insertion library into established communities. The library additionally enabled genome-wide measurement of gene-specific fitness during invasion, providing a functional readout of how community and dietary context affect the invading strain. Concurrent with invasion, half of the mice were switched from SD to a microbiota-accessible carbohydrate (MAC)-deficient high-sugar diet (MD), while the remaining mice were maintained on SD. After 28 days, all mice received a five-day course of ciprofloxacin to examine how antibiotic perturbation further affected community and MGE dynamics. Dense longitudinal stool sampling over 60 days enabled simultaneous tracking of community composition, invasion dynamics, and MGE activity (**Fig. 1A**).

**Figure 1:**
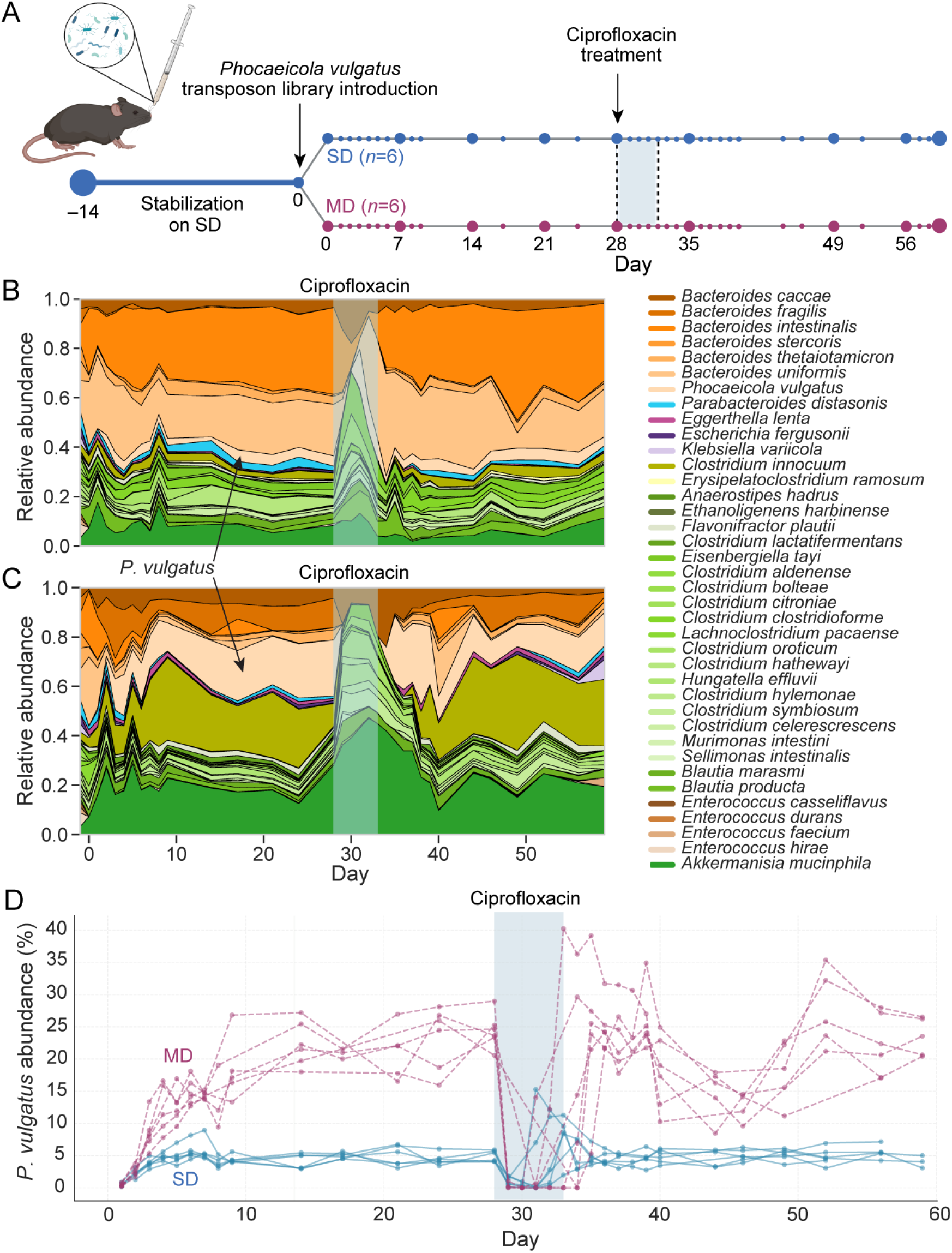
A synthetic gut community generates reproducible diet-dependent ecological states. A) Experimental design. Germ-free mice were colonized with a defined 38-member synthetic human gut community and maintained on standard diet (SD) for 14 days to allow community stabilization. At day 0, mice were separated into two dietary groups (*n*=6 per group) and gavaged with a barcoded transposon insertion library of *Phocaeicola vulgatus*. One group remained on SD, whereas the other was switched to a microbiota-accessible carbohydrate (MAC)-deficient high-sugar diet (MD). All mice received ciprofloxacin on days 28–32. Fecal samples were collected longitudinally at the indicated time points. B,C) Community composition of representative mice maintained on SD (B) or switched to MD (C) across the experimental time course. Relative abundances of the 38 community members were determined by metagenomic sequencing. MD induced substantial restructuring of the resident community, including reduced abundance of several Bacteroidota and expansion of *Akkermansia muciniphila* and members of Erysipelotrichaceae. Shading indicates the ciprofloxacin treatment period. D) Relative abundance of *P. vulgatus* across all mice. Each line represents an individual mouse. *P. vulgatus* remained at relatively low abundance in SD-fed mice, consistent with strong colonization resistance, but reached substantially higher relative abundance in MD-fed mice, establishing invasion-resistant and invasion-permissive ecological states, respectively.

This design generated two reproducible ecological contexts. Under SD conditions, *P. vulgatus* colonized all mice but remained at low abundance (4.4±0.9% on days 4, 14, and 24; **Fig. 1B,D**), indicating strong colonization resistance by the resident community. In contrast, MD-fed mice underwent rapid community restructuring (**Fig. 1C, S1**), including depletion of polysaccharide-degrading Bacteroidaceae and expansion of mucin-associated taxa, including *Akkermansia muciniphila*, as well as pathobiont-associated Erysipelotrichaceae. This restructuring is consistent with a shift from dietary polysaccharide utilization toward host mucin metabolism, while the contrasting invasion outcomes suggest that the dietary perturbation weakened the community’s barrier to invasion (*21*). Accordingly, *P. vulgatus* reached substantially higher abundance in MD-fed mice than in SD-fed mice (19.7±6.3% on days 4, 14, and 24; **Fig. 1C,D**). Ciprofloxacin treatment transiently reduced *P. vulgatus* abundance in MD-fed mice, but the species rebounded following treatment, whereas its abundance remained low in SD-fed mice. The dietary shift also led to the rapid loss of several resident taxa, including *Bacteroides intestinalis*, which persisted in SD-fed mice but declined below detection in MD-fed mice. Together, these contrasting ecological responses established invasion-resistant and invasion-permissive community states, providing a controlled framework to examine how ecological context influences horizontal gene transfer.

### Junction-resolved metagenomics reveals active and transferred MGEs

To track MGE dynamics in complex communities from short-read metagenomics data, we developed a junction-resolved framework that distinguishes integrated chromosomal forms from excised circular episomes (**Fig. 2A**). For integrative elements such as prophages and conjugative elements, excision generates episomal junctions absent from the chromosome, whereas integration into recipient genomes creates new host-element junctions (*6, 22*). By mapping paired-end reads spanning these diagnostic junctions and normalizing to host genome coverage, we quantified activation, defined as episome formation, and transfer, defined as integration into new host genomes, for 130 integrative MGEs (iMGEs) that showed evidence of excision across the 38-species community (**Fig. S2, Table S1**).

**Figure 2:**
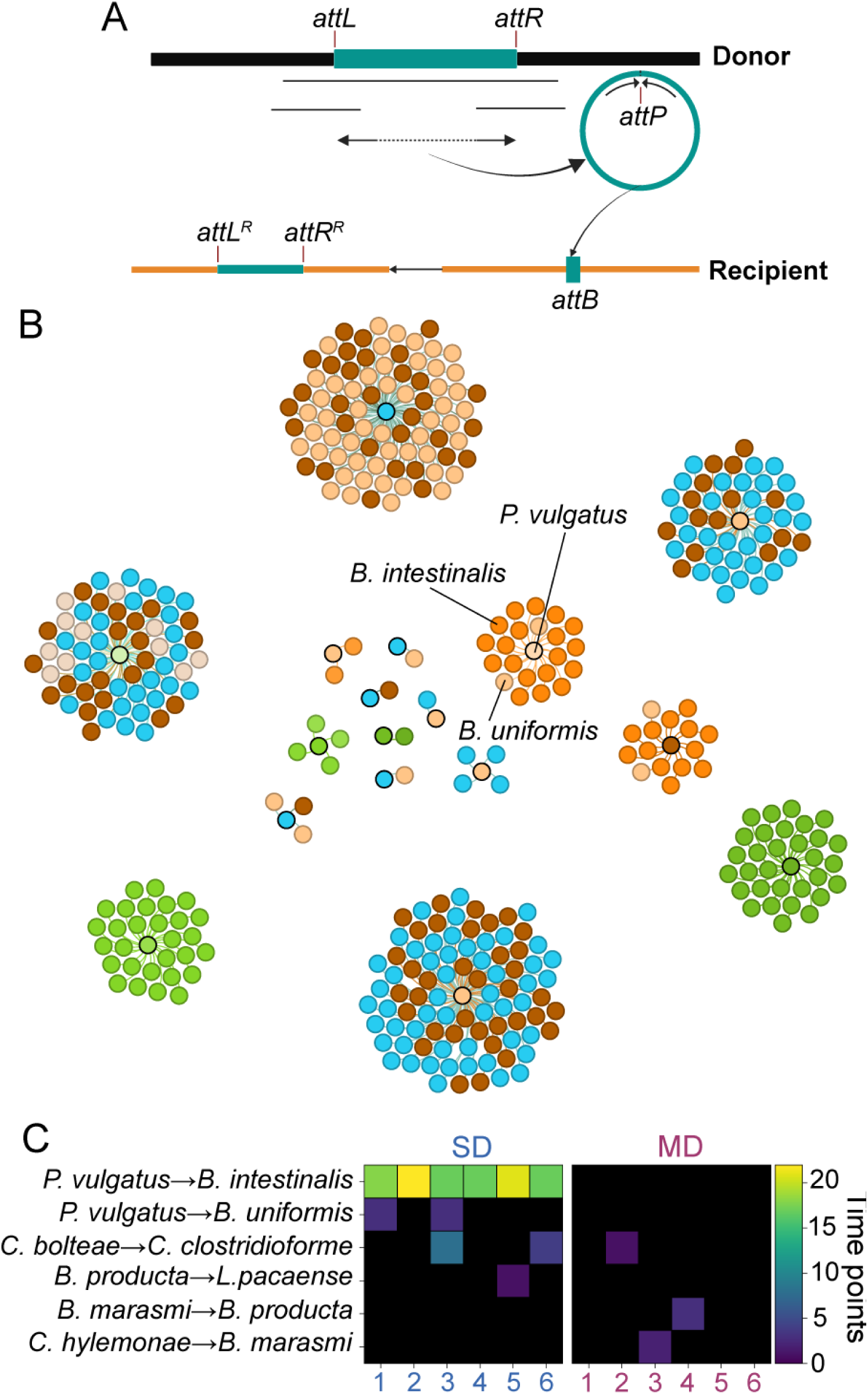
Junction-resolved metagenomics reveals rare horizontal transfer among active MGEs *in vivo*. A) Schematic of the junction-resolved strategy for detecting MGE excision and circularization from paired-end metagenomic sequencing. Excision of an integrated MGE generates a circular episome containing an episomal junction (*attP*) that is absent from the chromosomally integrated form. Reads spanning this diagnostic junction provide direct evidence of MGE excision and circularization. Integration into a recipient genome generates new host-MGE junctions (*attL^R^* and *attR^R^*), enabling *de novo* transfer events to be distinguished from pre-existing MGE–host associations. B) MGE-host association network for a representative mouse. Each hub represents a distinct MGE (episome), and connected nodes indicate species in which that MGE was (integration). Nodes are colored by bacterial species as indicated in Fig. 1. Connections among multiple species reveal MGEs with evidence of historical or *de novo* host associations. Labeled lines indicate *de novo* transfer events detected during the experiment, including recurrent transfer of the *P. vulgatus* prophage to *B. intestinalis* and subsequent transfer to *B. uniformis*. C) *De novo* MGE transfer events detected across individual mice during the 60-day experiment. Rows indicate donor–recipient transfer events and columns indicate individual mice maintained on SD (left; *n*=6) or MD (right; *n*=6). Color intensity indicates the number of time points at which each transfer event was detected. Despite widespread MGE activation, detectable *de novo* transfer was rare. Transfer of the *P. vulgatus* prophage to *B. intestinalis* was detected repeatedly in all six SD-fed mice but was absent from MD-fed mice.

This approach revealed extensive evidence of MGE exchange within the community prior to the start of this experiment. Eighteen iMGEs were present in multiple host genomes, either as identical sequences consistent with recent transfer or as divergent variants indicative of older transfer events (**Fig. 2B, S3A,B, Table S1**). All 18 exhibited episome formation during the experiment, indicating that previously transferred elements can remain active and retain the potential for remobilization long after integration. Long-read sequencing (Methods) of two samples independently confirmed both episomes and alternative integration sites detected by the short-read framework (**Fig. S4**).

We next asked whether this framework could identify *de novo* transfer during the 60-day experiment. By comparing *in vivo* integration junctions to baseline synteny networks from the input isolate genomes (Methods), we detected four new transfer events among members of the resident community. Three involved prophages and one involved an integrative mobile element (IME), with all four events occurring between members of the class Clostridia (**Fig. 2C, S3C, Table S1**). Thus, despite widespread MGE activation, successful transfer among established community members was rare and phylogenetically restricted.

A distinct and particularly recurrent transfer event involved the introduced invader: a prophage originating from *P. vulgatus* (HMPREF1058_RS10450-RS10085) repeatedly integrated into *B. intestinalis* across multiple mice, with subsequent transfer to *Bacteroides uniformis* in a subset of animals (**Fig. 2B,C, Fig. S3C**). Despite greater *P. vulgatus* invasion under MD conditions, this recurrent transfer occurred exclusively in SD-fed mice and was absent in MD-fed mice. This diet-dependent transfer from an invading strain to resident community members provided a tractable system for dissecting how ecological context governs HGT *in vivo*.

### Prophage activation is associated with invasion resistance

In SD-fed mice, where *P. vulgatus* remained at low abundance, prophage activation (measured by *attP* abundance, indicative of excised circular prophages) increased sharply during early colonization (days 1–9) reaching 17- to 40-fold above baseline in individual mice (**Fig. 3A**). Elevated *attP* abundance persisted throughout the experiment, including after antibiotic perturbation (days 28–32). In contrast, MD-fed mice, in which *P. vulgatus* achieved high abundance, exhibited little detectable prophage activation, with *attP* signals remaining near the detection limit (**Fig. 3B**). Across all animals, peak *attP* abundance was significantly higher in SD-fed than MD-fed mice (median fold-change: 22.0 versus 0.28; Wilcoxon rank-sum test, *p*=0.005). Strikingly, MD-fed mice exhibited substantially less prophage activation despite *P. vulgatus* reaching approximately fourfold higher relative abundance than in SD-fed mice, indicating that prophage activation was associated with ecological resistance to invasion rather than the relative abundance of the invader.

**Figure 3:**
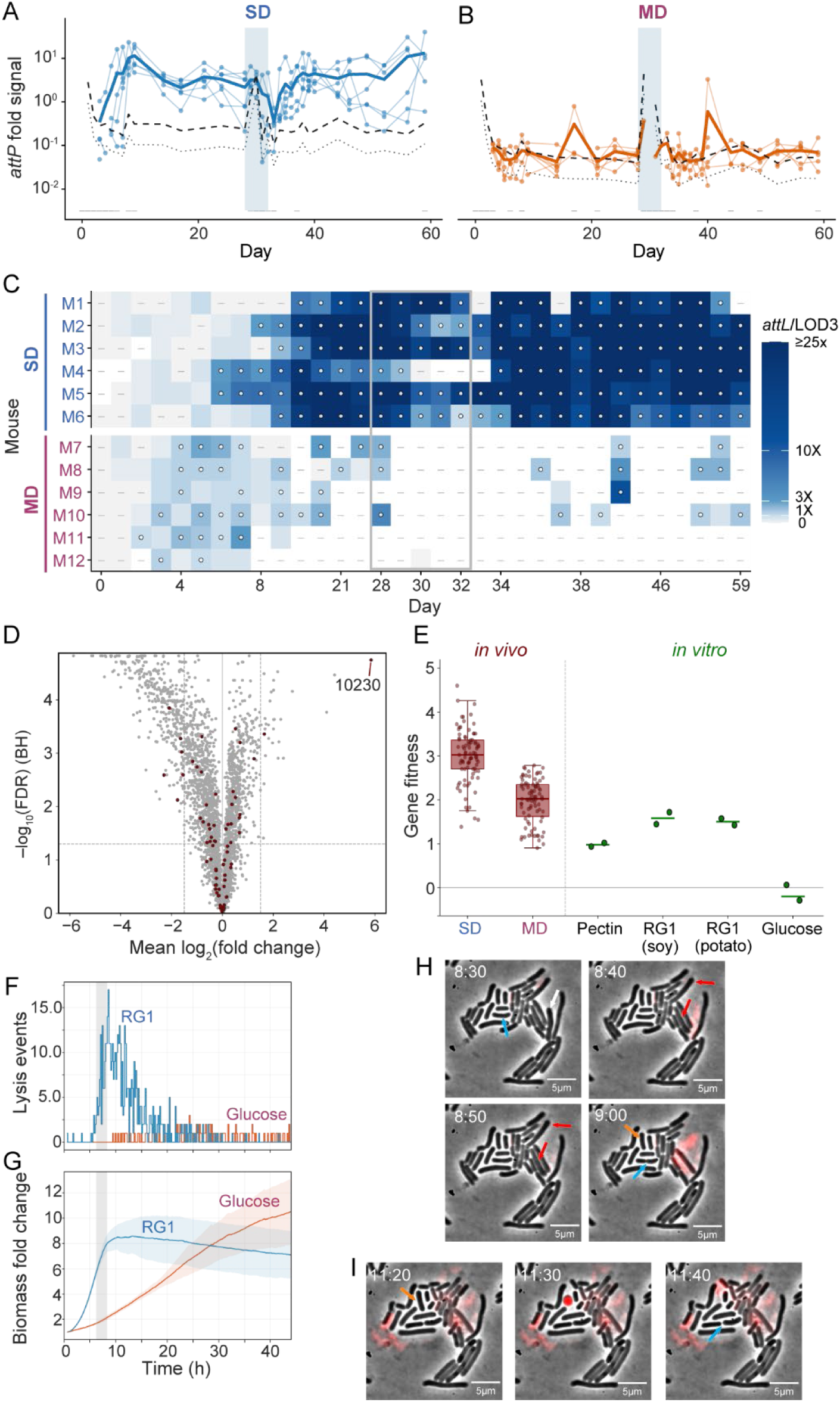
Dietary polysaccharides promote prophage activation, donor fitness costs, and lysis. A,B) Prophage activation dynamics in SD-fed (A) and MD-fed (B) mice. Prophage activation was quantified as the abundance of the episomal junction (*attP*) normalized to *P. vulgatus* genome coverage. Solid blue (SD) or orange (MD) lines indicate individual mice; dashed lines indicate three reads per *P.* vulgatus genome and dotted lines indicate one read per *P. vulgatus* genome. Prophage activation increased rapidly following invasion and remained elevated in SD-fed mice, whereas activation remained near or below the limit of detection in MD-fed mice despite the greater relative abundance of *P. vulgatus*. Shading indicates ciprofloxacin treatment. C) Longitudinal integration of the *P. vulgatus* prophage into *B. intestinalis*. Integration was quantified from the recipient-specific *attL* junction normalized to *B. intestinalis* genome coverage. Rows represent individual mice and columns represent sampling days; SD-fed mice (M1–M6) are shown above the horizontal line and MD-fed mice (M7–M12) below. Dots indicate samples in which *attL* abundance exceeded the detection threshold of three supporting reads (LOD3); dashes indicate samples with no read support. Integration emerged 5–10 days after invasion in all six SD-fed mice and subsequently persisted, whereas MD-fed mice showed only sporadic *attL* detection without sustained integration. D) Genome-wide fitness landscape of the *P. vulgatus* barcoded transposon library in SD-fed mice before antibiotic treatment. Mean log_2_ fold change in mutant abundance is plotted against statistical significance (−log_10_ FDR). Transferring prophage-associated genes are highlighted in red. Disruption of HMPREF1058_RS10230 (10230), encoding a prophage-associated terminase, conferred the largest fitness advantage detected genome-wide, indicating a substantial fitness cost associated with productive prophage activity. E) Fitness of HMPREF1058_RS10230 transposon mutants across *in vivo* and *in vitro* conditions. Points represent independent biological replicates or individual mouse samples from days 1–28 after colonization, and horizontal bars indicate mean fitness, with interquartile ranges for the *in vivo* conditions. Disruption of HMPREF1058_RS10230 conferred a strong fitness advantage *in vivo* and during growth on pectin and RG-I, but little or no advantage during growth on glucose. The stronger phenotype on RG-I than pectin is consistent with RG-I being a component of pectin and suggests that RG-I contributes to the carbon-source-dependent prophage phenotype. F,G) Single-cell lysis and population growth of *P. vulgatus* following transfer from rich medium to minimal medium containing soybean RG-I or glucose as the sole carbon source. (F) Timing of individual lysis events during time-lapse imaging. The dashed vertical line marks the time at which RG-I cultures began to depart from exponential growth and defines the subsequent interval over which lysis frequencies were compared between RG-I and glucose conditions. (G) Biomass accumulation during growth on RG-I or glucose. Solid lines indicate means across biological replicates and shaded regions indicate 1 standard deviation. *P. vulgatus* initially grew rapidly on RG-I, with biomass accumulation comparable to or exceeding that on glucose, but growth subsequently slowed as lysis increased after ∼3–4 generations. Lysis remained rare on glucose, resulting in substantially greater biomass accumulation at later time points than on RG-I. H,I) Representative time-lapse microscopy of *P. vulgatus* growing on soybean RG-I, illustrating distinct cellular trajectories preceding lysis. The blue arrow marks a cell that continues to grow and divide, whereas the white arrow marks a cell that undergoes abrupt lysis. Red arrows mark cells that cease growth before lysing. The orange arrow marks a cell that loses its characteristic rod-like morphology, becomes PI-positive, and subsequently lyses. Scale bar: 5 µm.

### Prophage activation precedes stable integration into resident populations

Having identified a recurrent prophage transfer event from the invading *P. vulgatus* strain to resident community members, we next resolved its temporal dynamics. In all six SD-fed mice, *attL* junctions corresponding to integration of the *P. vulgatus* prophage into *B. intestinalis* became detectable 5–10 days after invasion and remained elevated throughout the experiment (**Fig. 3C**). We inferred from single nucleotide polymorphism (SNP) profiles that transfer likely occurred independently in most mice, with *attL* signals rising to high and persistent levels (10^-1^–10^0^ *attL* junctions per *B. intestinalis* genome), indicating stable establishment of prophage-containing recipient populations rather than transient infection.

These integration dynamics followed the earlier rise in *attP* abundance (**Fig. 3A**), consistent with prophage activation preceding successful horizontal transfer. In contrast, MD-fed mice exhibited little evidence of sustained integration in *B. intestinalis* despite substantially greater invasion success by *P. vulgatus*. *attL* junctions remained near the limit of detection throughout the experiment, with only occasional transient signals that failed to persist (**Fig. 3C**). Long-read sequencing of selected SD samples confirmed integration of the transferred prophage at the predicted *B. intestinalis* attachment site (*attB*) and confirmed the identity of the transferred sequence (**Fig. S5**).

The absence of transfer in MD-fed mice reflected not only reduced prophage activation but also loss of the primary recipient population. Whereas *B. intestinalis* persisted throughout the experiment in SD-fed mice, it rapidly declined below undetectable levels following the diet switch in MD-fed mice (**Fig. S6**), coincident with expansion of *P. vulgatus* and blooms of Erysipelotrichaceae. Thus, community restructuring eliminated the dominant recipient population before stable transfer could be established. Notably, transfer was not redirected to other abundant *Bacteroides* species, including *B. uniformis*, suggesting that recipient permissiveness is species-specific.

Together, these results show that the striking diet dependence of prophage transfer reflected two effects of community context: prophage activation was strongly suppressed in MD-fed mice, and the primary recipient population was lost following dietary restructuring.

### Prophage activation imposes a fitness cost on the invading strain

To determine whether prophage activation affected invader fitness, we leveraged random barcode transposon insertion sequencing (RB-TnSeq) (*23*) of the *P. vulgatus* library throughout colonization. Genome-wide fitness analysis identified disruption of a gene in the transferring prophage locus, HMPREF1058_RS10230, which encodes a terminase protein, as the strongest beneficial mutation in both dietary contexts (mean log_2_(fold change) = 5.9, FDR 1.8×10^-5^; **Fig. 3D, S7**). Because terminase proteins are required for DNA packaging and productive virion assembly, this result suggests that disabling prophage production substantially increases fitness during gut colonization.

The magnitude of this fitness benefit depended on ecological context. Disruption of HMPREF1058_RS10230 conferred a larger advantage in SD-fed mice, where prophage activation was sustained (**Fig. 3A, S7B**), than in MD-fed mice, where activation remained suppressed (**Fig. 3B, S7B**). The correspondence between prophage activation and the fitness benefit of terminase disruption indicates that active virion production imposes a substantial cost on *P. vulgatus* during gut colonization.

### Diversity-generating retroelements are associated with host-range expansion during transfer

To investigate mechanisms that might facilitate transfer into resident species, we examined sequence variation within the transferring prophage. Single-nucleotide variant analysis identified seven mutational hotspots, including two that repeatedly accumulated mutations in every SD-fed mouse: a receptor-binding protein (RBP, HMPREF1058_RS10405) and a ParB-Related, ThiF-Related Cassette (PRTRC) system protein B (HMPREF1058_RS10145; **Fig. 4A-D**). The RBP is adjacent to a diversity-generating retroelement (DGR) (*24*), a targeted mutagenesis system that uses error-prone reverse transcription of a template sequence to repeatedly diversify a corresponding variable region with the RBP gene and PRTRC protein B. Longitudinal haplotype analysis identified 20 RBP variants whose frequencies increased concurrently with integration into *B. intestinalis* across all six mice (29 haplotype-mouse instances; **Fig. 4E,S8**). These variants were significantly enriched during integration windows relative to neutral expectations (permutation test, *p*<0.0001), with individual haplotypes increasing in frequency by 0.15–0.60. The variable residues mapped to a predicted C-type lectin domain (CLec3) (*25*) (**Fig. 4D**), a structural class commonly targeted by DGR systems enabling extensive sequence variation (*26, 27*), suggesting that DGR-mediated diversification may facilitate prophage host-range expansion during transfer. Notably, DGR-mediated diversification was largely absent in MD-fed mice, where successful transfer was also not observed (**Fig. 4B,C**).

**Figure 4:**
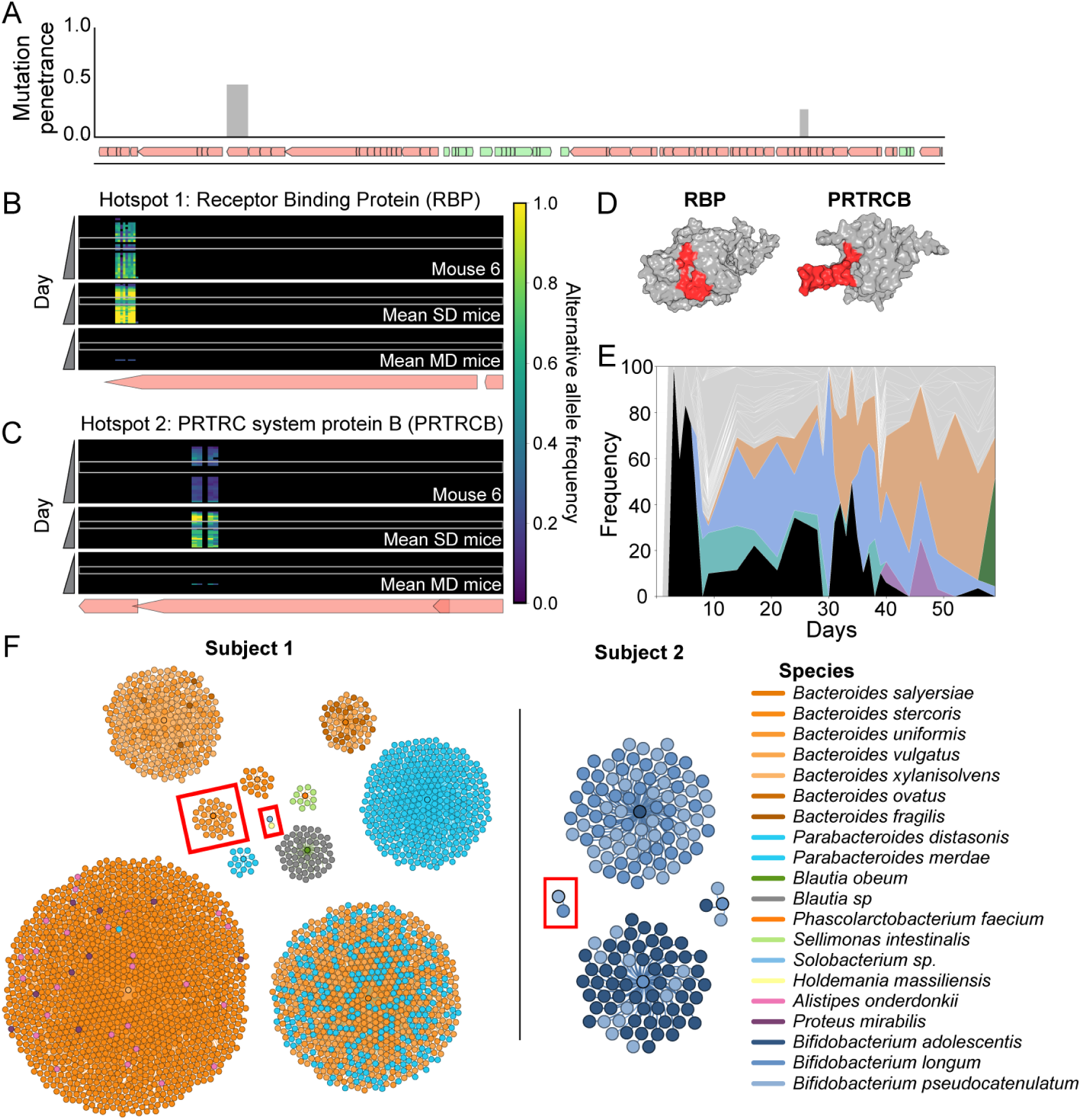
Junction-resolved metagenomics reveals prophage diversification and horizontal transfer in mice and humans. A) Distribution of sequence variation across the *P. vulgatus* prophage genome. The gene map is shown below, with genes colored by strand orientation. Bars indicate the fraction of samples containing sequence variation at each genomic position. Two recurrent regions of variation were identified, including prominent variation within a receptor-binding protein (RBP; hotspot 1) and protein B of a PRTRC system (hotspot 2). B,C) Longitudinal sequence variation within the prophage RBP (B) and PRTRC system protein B (C). Alternative allele frequencies are shown across genomic position and time. For each locus, the top panel shows a representative SD-fed mouse (M6), the middle panel shows the mean across SD-fed mice, and the bottom panel shows the mean across MD-fed mice. Sequence variation increased reproducibly in SD-fed mice but remained limited in MD-fed mice. Gray shading indicates ciprofloxacin treatment. D) AlphaFold2-predicted structure of the prophage RBP (HMPREF1058_RS10405) and PRTRC system protein B (HMPREF1058_RS10145). Residues exhibiting recurrent sequence variation are highlighted in red; these residues in the RBP cluster within a predicted C-type lectin-like domain implicated in ligand and glycan recognition. E) Stacked-area plot showing the frequency dynamics of RBP haplotypes identified in mouse 6. All haplotypes reaching ≥10% and showing increasing frequency from beginning to end are shown individually, with rare variants grouped in gray. F) Application of junction-resolved metagenomics to longitudinal human gut metagenomes from the BIO-ML cohort (*29*). MGE–host association networks are shown for two representative participants. Nodes represent bacterial species and edges indicate MGE–host associations detected across longitudinal samples. Red boxes highlight candidate *de novo* transfer events that emerged during the sampling period, illustrating the application of junction-resolved analysis to retrospectively identify interspecies MGE transfer in longitudinally sampled human microbiomes.

### Rhamnogalacturonan I induces cell lysis consistent with prophage activation

Our *in vivo* RB-TnSeq data indicated that productive prophage activity imposes a substantial fitness cost on *P. vulgatus* under SD conditions, raising the possibility that components of the polysaccharide-rich diet stimulate prophage production. To identify potential dietary triggers, we examined the fitness of the *P. vulgatus* transposon library during growth in defined minimal media containing individual carbon sources. Disruption of HMPREF1058_RS10230 conferred no detectable fitness advantage during growth on glucose but conferred a positive fitness effect during growth on citrus pectin (fitness = 1.9), in the same direction as the strong advantage observed in SD-fed mice (**Fig. 3D,E**). We therefore tested rhamnogalacturonan I (RG-I), a major structural component of pectin, and found that the fitness advantage increased further during growth on potato- and soybean-derived RG-I (fitness = 3.0 and 3.1, respectively; **Fig. 3E**). Thus, RG-I recapitulated a substantial component of the prophage-associated fitness cost observed *in vivo*.

We therefore directly tested whether the contrasting fitness phenotypes in glucose and RG-I were accompanied by differences in cell death consistent with productive prophage induction. *P. vulgatus* cells grown in rich medium were washed and transferred to minimal medium containing either glucose or soybean-derived RG-I, and individual cells were followed by anaerobic time-lapse microscopy (Methods). Cells growing on glucose continued dividing for ∼7 generations, with lysis observed in only 0.1% of cells (**Fig. 3F,G; Movie S1**). In RG-I, *P. vulgatus* initially grew more rapidly than in glucose for ∼3 generations, demonstrating efficient utilization of the polysaccharide, before a subset abruptly stopped growing and lysed within 10–30 min; other cells first lost their characteristic rod-like morphology, followed by propidium iodide entry and eventual lysis(**Fig. 3F-I; Movie S2**). Overall, 5.8% of cells underwent lysis during growth on RG-1, a >50-fold increase relative to glucose.

The delayed onset of lysis after several generations of growth argues against an immediate toxic effect of RG-I and instead supports activation of a lytic process during growth on this polysaccharide (*28*). Together with the carbon-source-dependent fitness advantage conferred by disruption of the transferring prophage terminase gene required for virion production, these observations support a model in which growth on RG-I promotes productive prophage activity in *P. vulgatus*, resulting in loss of cellular integrity and lysis. These findings provide a direct mechanistic link between dietary polysaccharides enriched under the invasion-resistant condition, the fitness cost of productive prophage activity, and the prophage activation observed *in vivo*.

### Junction-resolved metagenomics detects HGT in longitudinal human microbiomes

To test whether our junction-resolved framework could detect HGT in existing longitudinal human microbiome datasets, we analyzed publicly available metagenomes from three densely sampled healthy individuals (*29*). Because these samples were collected without experimental manipulation or selection for transferred elements, this analysis provided an opportunity to test whether MGE transfer could be resolved retrospectively from naturally sampled microbiomes.

Across the three individuals, we detected three candidate *de novo* transfer events in two of the individuals, including two prophages and one IME (**Fig. 4J**). These comprised prophage transfer from *Bacteroides stercoris* to *Bacteroides uniformis*, prophage transfer from *Bifidobacterium pseudocatenulatum* to *Bifidobacterium longum*, and IME transfer from a *Solobacterium* species to *Ruthenibacterium lactatiformans*. In each case, the recipient-specific MGE-host association emerged during longitudinal sampling. The prophage transfer from *B. stercoris* to *B. uniformis* persisted for 26 time points, suggesting a fitness advantage in the recipient.

These findings demonstrate that junction-resolved metagenomics can be applied retroactively to existing longitudinal human metagenomic datasets to identify candidate HGT events, extending the framework beyond experimentally controlled communities and providing a means to investigate MGE dynamics in naturally sampled microbiomes.

## Discussion

By combining dense longitudinal sampling with junction-resolved metagenomic analysis, we tracked MGE activation and HGT in a complex mammalian gut community, distinguishing historical transfer events from *de novo* transfer occurring during the experiment. A striking observation was the disconnect between activation and successful transfer. Although 130 MGEs formed detectable episomes, only six *de novo* transfer events were detected over 60 days. These findings suggest that activation is common, whereas successful establishment in a new host is rare, indicating that the principal barrier to HGT may not be MGE mobilization itself but rather ecological and evolutionary constraints on transfer and persistence.

Among the few successful transfer events, one dominated the landscape: a prophage from an invading *P. vulgatus* strain repeatedly integrated into resident *B. intestinalis* populations in every mouse maintained on a standard polysaccharide-rich diet, with subsequent transfer to *B. uniformis* in a subset of animals. Paradoxically, transfer occurred exclusively under conditions that resisted invasion. In SD-fed mice, where resident communities maintained strong colonization resistance and *P. vulgatus* remained at low abundance, prophage activation was elevated 20- to 40-fold and transfer was robust and reproducible. In contrast, in MD-fed mice, where community restructuring permitted *P. vulgatus* expansion, prophage activation remained low and transfer was undetectable. These observations challenge the intuitive expectation that transfer opportunities scale with donor abundance. Instead, they suggest that ecological interactions imposed by the resident community can stimulate prophage activation and thereby determine when donor cells become sources of horizontal transfer.

Successful transfer required alignment of multiple ecological conditions. In SD-fed mice, prophage activation coincided with persistence of *B. intestinalis*, the dominant recipient population. In MD-fed mice, transfer failed not only because prophage activation was suppressed but also because *B. intestinalis* rapidly declined below detection following the dietary shift. Notably, transfer was not redirected to *B. uniformis*, which remained detectable in MD-fed mice, suggesting that recipient permissiveness is species-specific and that successful transfer to *B. uniformis* may depend on prior evolution within *B. intestinalis*. Together, these observations indicate that HGT emerges from the intersection of donor activation, ecological opportunity, and recipient compatibility.

The temporal resolution afforded by our approach revealed evolutionary dynamics accompanying this transfer process. Prophage activation preceded integration into *B. intestinalis* by several days, during which DGR activity diversified the prophage RBP. Twenty RBP haplotypes increased in frequency during transfer, with individual variants rising by as much as 60% across independent mice. These variants mapped to a C-type lectin domain commonly targeted by DGRs to accommodate extensive sequence variation, providing *in vivo* support for the hypothesis that prophage-associated DGRs can facilitate expansion into new recipient hosts. Rather than generating diversity in anticipation of future ecological opportunities, diversification occurred during an active transfer window and was tightly coupled to emergence in a novel recipient species.

Prophage activity also imposed a substantial fitness cost on the invading strain, revealing how dietary substrates can regulate the donor side of the transfer process. Disruption of a prophage terminase gene required for production of functional virions conferred the largest fitness benefit detected genome-wide, with stronger effects in SD-fed mice where activation was sustained. *In vitro*, this fitness advantage was carbon-source dependent: terminase disruption had little effect during growth on glucose, a substantial effect on pectin, and an even greater effect on RG-I. This genetic signature was accompanied by RG-I-specific explosive cell lysis that was not observed during growth on glucose, providing direct phenotypic support for prophage-mediated killing. Together, these observations link the dietary environment to prophage-associated killing and its fitness consequences for the donor.

Notably, strong positive selection was observed against this virion-production function but not against other prophage functions. This pattern raises the possibility that retention of the prophage provides benefits to the bacterial host even though productive phage formation and the associated risk of lysis are costly. Activation and transfer occurred before antibiotic perturbation in all mice, demonstrating that ecologically relevant environmental signals—not antibiotics or direct selection for transferred genes—can drive substantial gene flow in complex communities.

The strong disparity between activation and successful transfer suggests that compatible recipients, rather than MGE activation itself, are often the limiting factor for HGT. Although 130 MGEs exhibited episome formation, only six *de novo* transfer events were detected. These constraints likely include compatibility of integration and replication machinery (*30, 31*), evasion of host defenses (*32*), and recognition of appropriate surface receptors (*33, 34*). Additional barriers may arise from competition among related MGEs. For example, a homologous prophage in *B. stercoris* occupies the same tRNA integration site targeted by the *P. vulgatus* prophage, potentially preventing transfer through site exclusion. More generally, the long history of coevolution within this donor-derived community may have reduced opportunities for successful transfer compared with communities undergoing active assembly, disturbance, or invasion.

A key advantage of our approach is that it utilizes standard short-read metagenomic sequencing, making it broadly accessible and readily applicable to existing longitudinal datasets. By tracking diagnostic junctions, we resolved activation, transfer, diversification, and recipient population dynamics over time without requiring cultivation, engineered markers, or routine long-read sequencing. Importantly, this framework enables experimental tracking of HGT without selecting for antibiotic resistance, metabolic traits, or the transferred element itself, avoiding perturbations that can obscure the ecological drivers of gene transfer. Although long-read approaches remain valuable for structural validation, junction-resolved metagenomics provides a scalable framework for studying MGE dynamics at temporal resolutions that are difficult to achieve with current long-read, proximity-ligation, or single-cell approaches. We further applied the framework to a longitudinal human metagenomic dataset, demonstrating its ability to retrospectively resolve transfer in naturally sampled microbiomes and highlighting its potential for systematic studies of HGT across diverse microbial ecosystems, from host-associated microbiomes to environmental communities.

Together, our findings suggest that HGT is not simply a consequence of genetic compatibility between mobile elements and their hosts. Instead, transfer emerges from ecological interactions that determine when donors activate, which recipients persist, and whether newly transferred elements can establish within recipient populations. Diet can influence multiple steps in this process, from prophage activation and donor lysis to recipient availability, while diversification accompanies transfer between populations. By resolving these processes in real time, our work reveals how environmental factors such as diet shape the movement of genes through microbial communities and provides a framework for understanding how ecological change influences microbial evolution across diverse ecosystems.

## Figures

**Figure S1:**
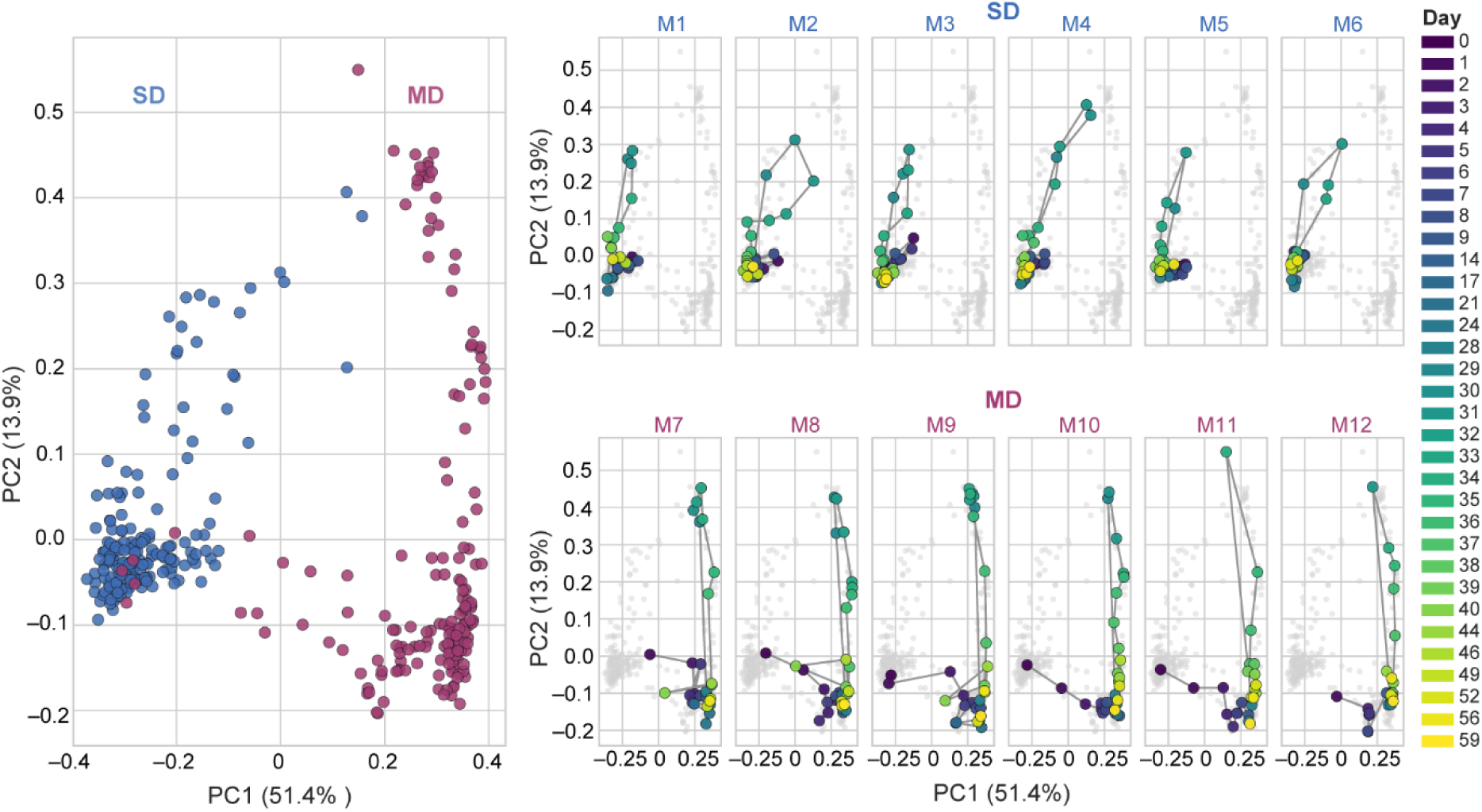
Diet drives gut microbial community composition, with reproducible temporal trajectories across individual mice. Principal coordinates analysis (PCoA) of Bray-Curtis dissimilarities calculated from species-level relative abundances across all longitudinal fecal samples. Axes show principal coordinates 1 and 2, with the percentage of total variation explained by each axis indicated in parentheses. Left: ordination of all samples colored by diet, showing clear separation of standard diet (SD)- and MAC-deficient diet (MD)-fed communities along PC1 (SD, blue; MD, red). Right: the same ordination shown separately for each mouse (M1–M12; top row, SD-fed mice M1–M6; bottom row, MD-fed mice M7–M12). Gray points show all samples for reference, and colored points show samples from the indicated mouse, connected in temporal order to depict its composition trajectory. Point color indicates sampling day (dark purple, early; yellow, late), as indicated by the shared color bar. Individual mice followed reproducible diet-dependent trajectories, with additional displacement and subsequent recovery associated with ciprofloxacin treatment.

**Figure S2:**
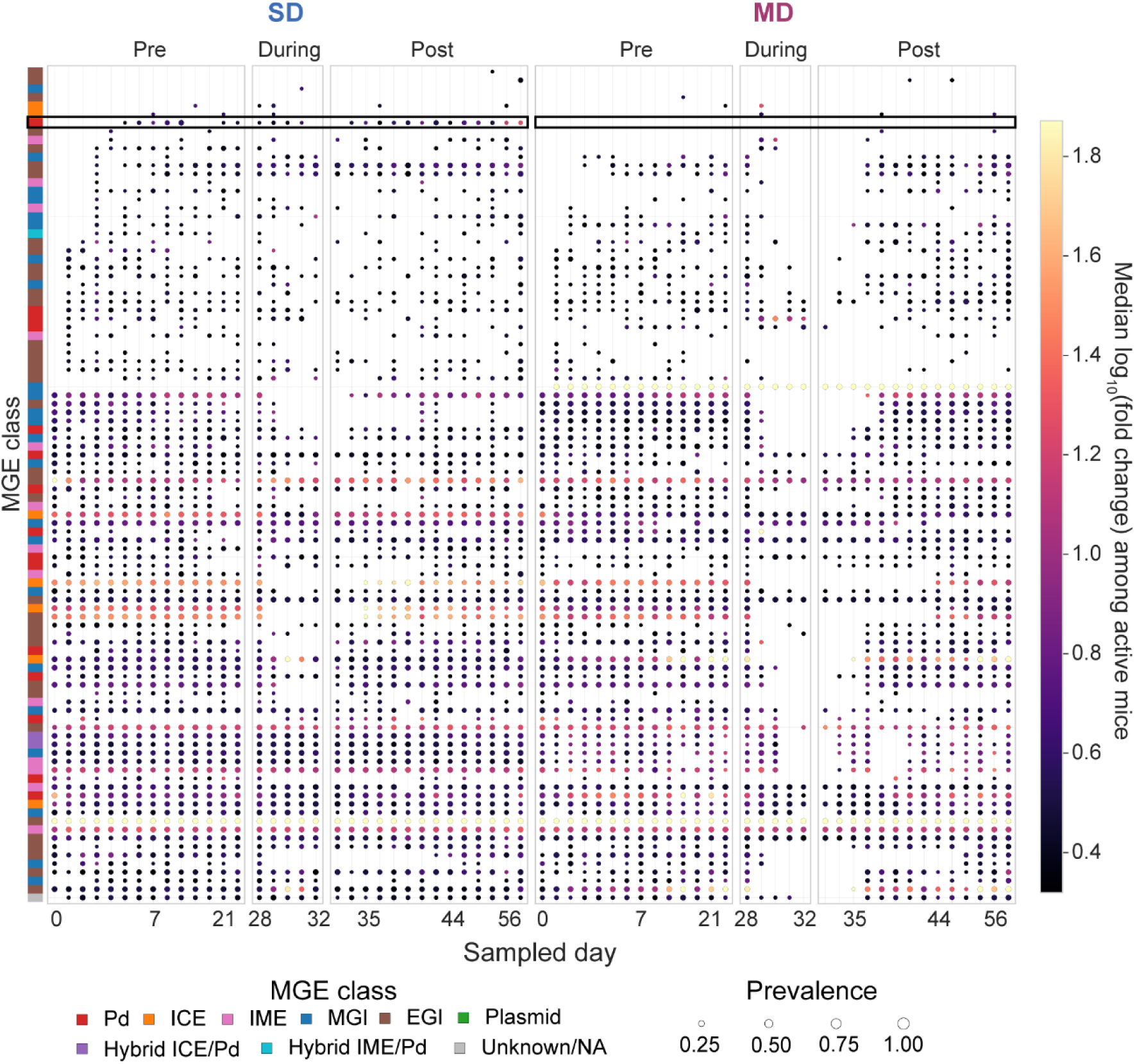
Widespread MGE activation exhibits element- and diet-dependent dynamics. Bubble heatmaps show longitudinal episome formation across circularizing MGEs detected in the synthetic gut community. Each row represents a distinct MGE, and columns represent sampling days. MGEs are ordered identically for both diets (**Table S1**), according to their detected appearance in either diet. Bubble size indicates the fraction of mice in which the corresponding episomal junction was detected at each time point, and bubble color indicates the median log_10_-transformed episomal junction abundance per host genome equivalent among mice in which the junction was detected. SD-fed mice are shown at left and MD-fed mice at right. Episome formation was widespread but heterogeneous, revealing strong element-specific and diet-dependent differences in MGE activation. The highlighted region indicates the transferring *P. vulgatus* prophage. The color bar at left indicates MGE classification: Pd, prophage-derived; ICE, integrative and conjugative element; IME, integrative mobilizable element; MGI, mobile genetic island containing an identifiable recombinase; EGI, excisable genomic island lacking an obvious recombinase or mobilization genes.

**Figure S3:**
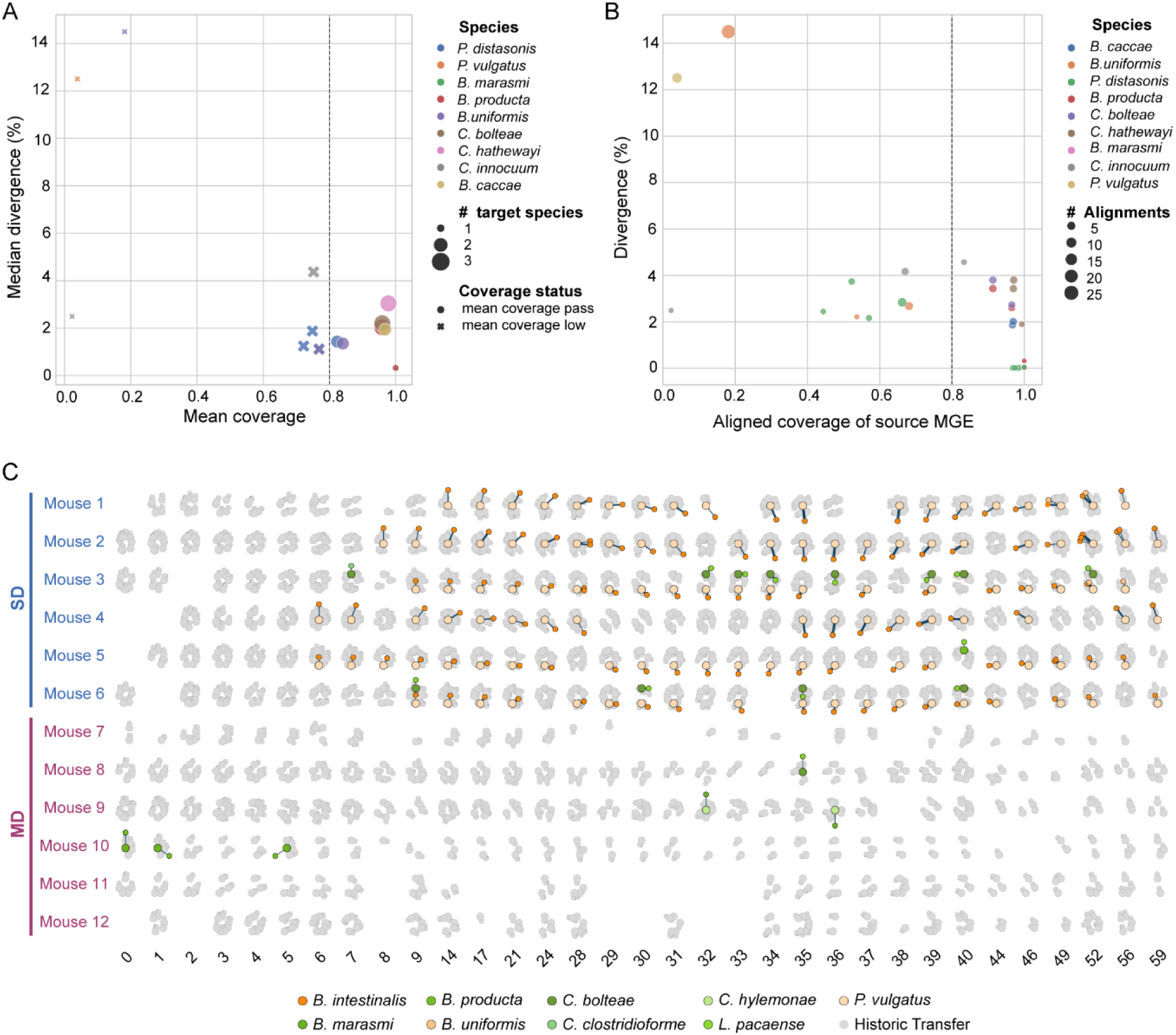
Filtering distinguishes *de novo* MGE transfer from pre-existing genomic similarity. A) Sequence divergence and aligned coverage between source MGEs and candidate recipient genomes. Each point represents a source MGE–recipient species pair, colored by source species; point size indicates the number of candidate recipient species associated with the MGE. Filled circles indicate comparisons with mean aligned coverage ≥0.8, whereas crosses indicate lower-coverage comparisons. The dashed vertical line marks the 0.8 aligned-coverage threshold. High-coverage, low-divergence comparisons are consistent with closely related MGE sequences shared among genomes, whereas lower-coverage or more divergent comparisons distinguish partial homology from near-identical MGE sequences. B) Sequence divergence and aligned coverage for individual source MGE–recipient genome alignments. Each point represents an individual alignment, colored by source species, with point size indicating the number of alignments contributing to the comparison. The dashed vertical line indicates the 0.8 aligned-coverage threshold used to distinguish broadly conserved MGE sequence from more limited sequence homology. The distribution of coverage and divergence illustrates the range of sequence similarity underlying candidate MGE–host associations and informed filtering of events attributable to pre-existing homology rather than *de novo* transfer. C) Longitudinal MGE–host association networks before and after filtering for *de novo* transfer. Rows represent individual mice and columns represent sampling days. Gray networks show all detected MGE–host associations, whereas colored nodes and edges indicate associations passing the criteria for *de novo* transfer. Large colored nodes denote MGE source species and smaller colored nodes denote recipient species; node colors indicate bacterial species shown at bottom. Candidate *de novo* transfers were required to represent high-confidence junction-resolved integration events that were absent from the baseline community and passed sequence-homology and recurrence filters. SD-fed mice are shown in rows 1–6 and MD-fed mice in rows 7–12.

**Figure S4:**
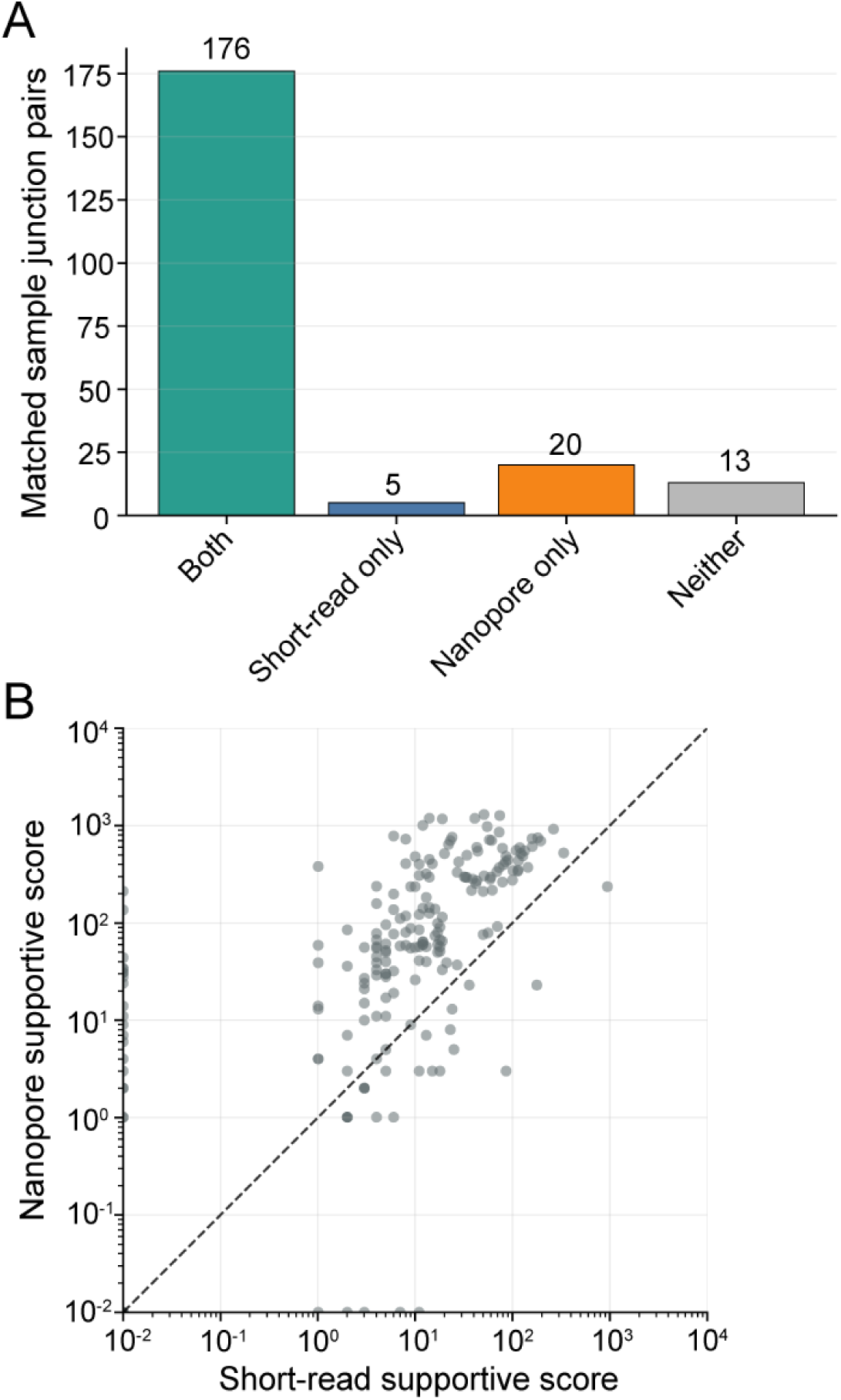
Cross-platform validation of MGE junction detection. A) Concordance of MGE junction detection between short-read and Oxford Nanopore sequencing of matched metagenomic samples. Bars indicate the number of MGE junctions evaluated across two matched samples that were detected by both platforms, by short-read sequencing only, by Nanopore sequencing only, or by neither platform at these time points. Of 214 junctions evaluated, 176 (82%) were detected by both platforms, demonstrating broad cross-platform concordance of junction detection. B) Comparison of sequencing support for MGE junctions detected by both platforms. Each point represents a junction detected in a matched sample. The dashed line indicates equal support from the two platforms. Junction support was quantified using a support score combining directly junction-spanning reads with a weighted contribution from local bridge reads. Support scores were strongly correlated in rank across platforms but showed a weaker linear relationship (Spearman *ρ*=0.73; Pearson *r*=0.31).

**Figure S5:**
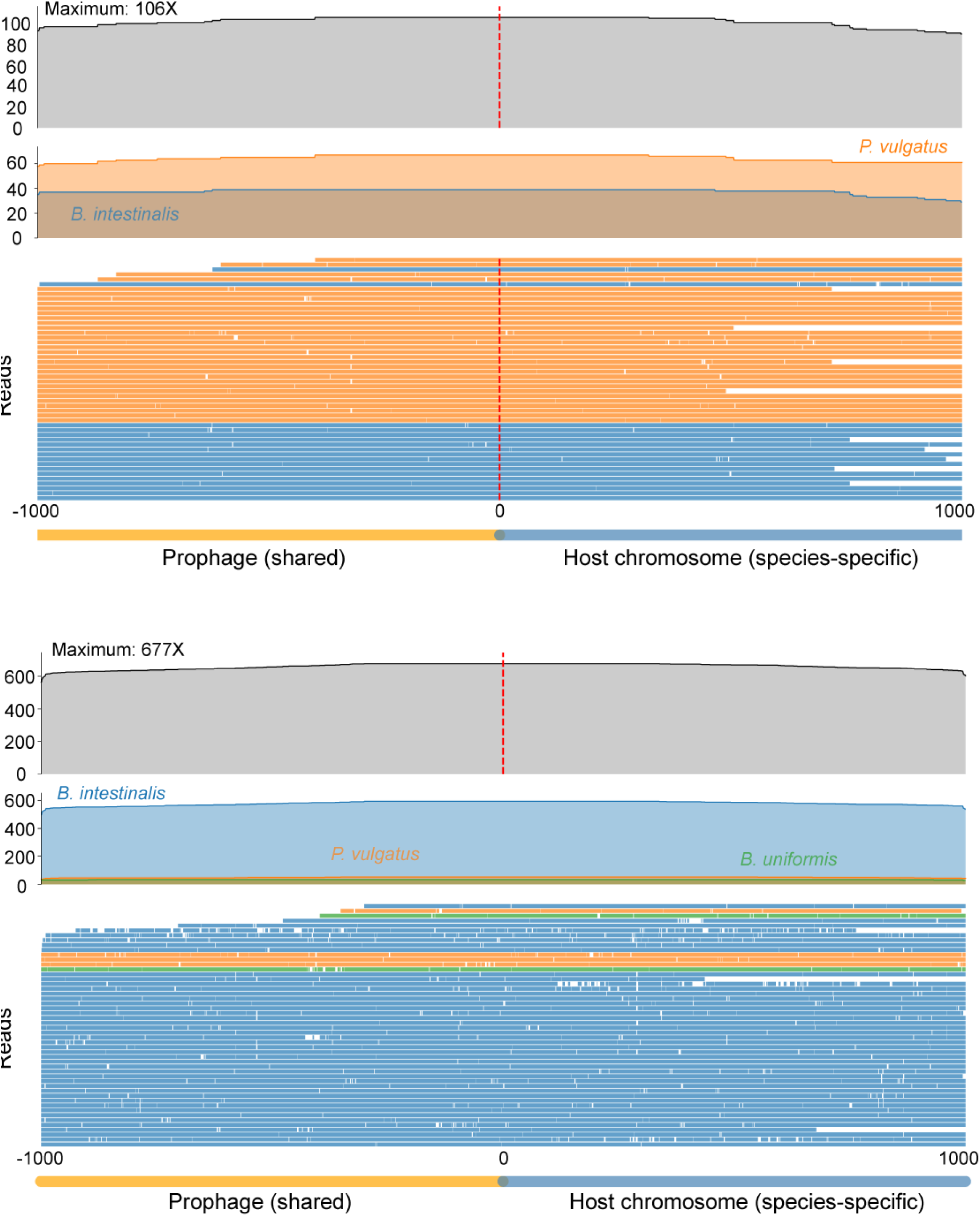
Long-read sequencing validates sequential transfer of the *P. vulgatus* prophage across recipient species. A) Nanopore read support for integration of the *P. vulgatus* prophage into *B. intestinalis* on day 8 in mouse 2. The top track shows total sequencing coverage across the integration site, and the middle track shows coverage assigned to individual species. The bottom panel shows a subsampled set of individual Nanopore read alignments spanning the prophage–host junction (red dashed line). Reads extending continuously from *P. vulgatus* prophage sequence into *B. intestinalis*-specific chromosomal sequence provide direct long-read support for integration of the prophage into the recipient genome. B) Nanopore read support for integration of the *P. vulgatus* prophage on day 52 in mouse 3. Tracks are displayed as in (A). Individual long reads spanning from prophage sequence into *B. intestinalis*- or *B. uniformis*-specific chromosomal sequence directly validate the secondary recipient identified by junction-resolved short-read metagenomics.

**Figure S6:**
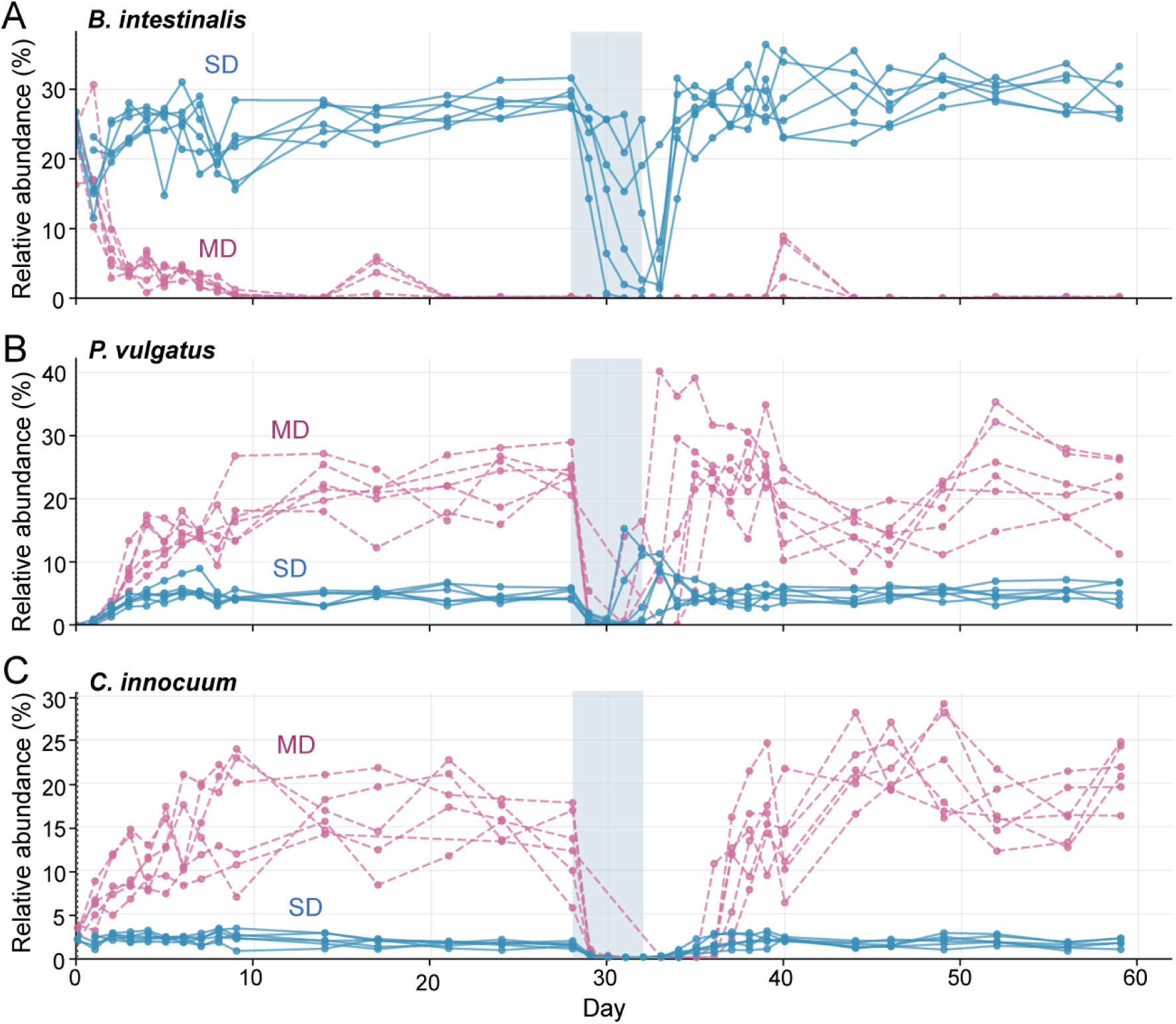
Diet-dependent abundance dynamics of *B. intestinalis*, *P. vulgatus*, and *C. innocuum*. Relative abundance trajectories of selected taxa across the 60-day experiment. Each line represents an individual mouse maintained on SD (blue) or MD (pink). Shading indicates ciprofloxacin treatment (days 28–32). A) *B. intestinalis*, the primary recipient of the *P. vulgatus* prophage, persisted at high relative abundance in SD-fed mice but rapidly declined following the switch to MD and remained below or near the limit of detection (≥0.5%) for most of the experiment. *B. intestinalis* abundance transiently decreased during ciprofloxacin treatment in SD-fed mice and subsequently recovered. B) *P. vulgatus* remained at relatively low abundance following invasion of SD-fed communities but reached substantially higher relative abundance in MD-fed mice. Ciprofloxacin treatment transiently reduced *P. vulgatus* abundance in both dietary groups, followed by recovery. C) *C. innocuum* expanded following the switch to MD and remained at substantially higher relative abundance in MD-fed than SD-fed mice throughout most of the experiment. Ciprofloxacin treatment resulted in a transient decrease followed by recovery.

**Figure S7:**
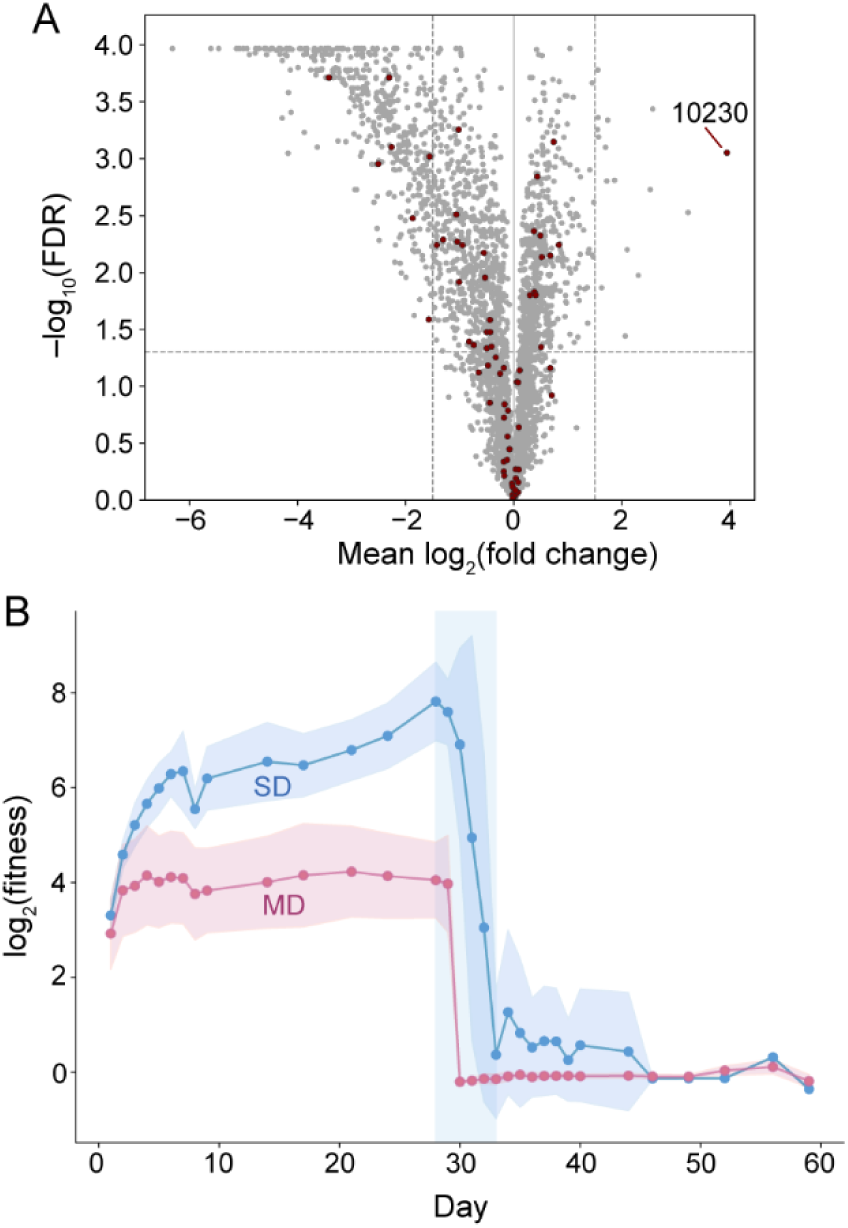
Disruption of the prophage terminase confers a diet-dependent fitness advantage *in vivo*. A) Genome-wide fitness landscape of the *P. vulgatus* barcoded transposon library in MD-fed mice before antibiotic treatment. Mean log_2_ fold change in mutant abundance is plotted against statistical significance (−log_10_ FDR). Prophage-associated genes are highlighted in red. Disruption of HMPREF1058_RS10230 (10230), encoding a prophage-associated terminase, conferred the largest fitness advantage detected genome-wide, indicating a substantial fitness cost associated with terminase function *in vivo*. B) Longitudinal fitness of HMPREF1058_RS10230 transposon mutants in SD- and MD-fed mice. Lines indicate mean fitness across mice and shaded regions indicate SD. The fitness advantage of terminase disruption increased progressively in SD-fed mice before antibiotic treatment, whereas the advantage was smaller and remained relatively stable in MD-fed mice. Following ciprofloxacin treatment, the fitness advantage rapidly declined in both dietary conditions. Shaded rectangle indicates the ciprofloxacin treatment period.

**Figure S8:**
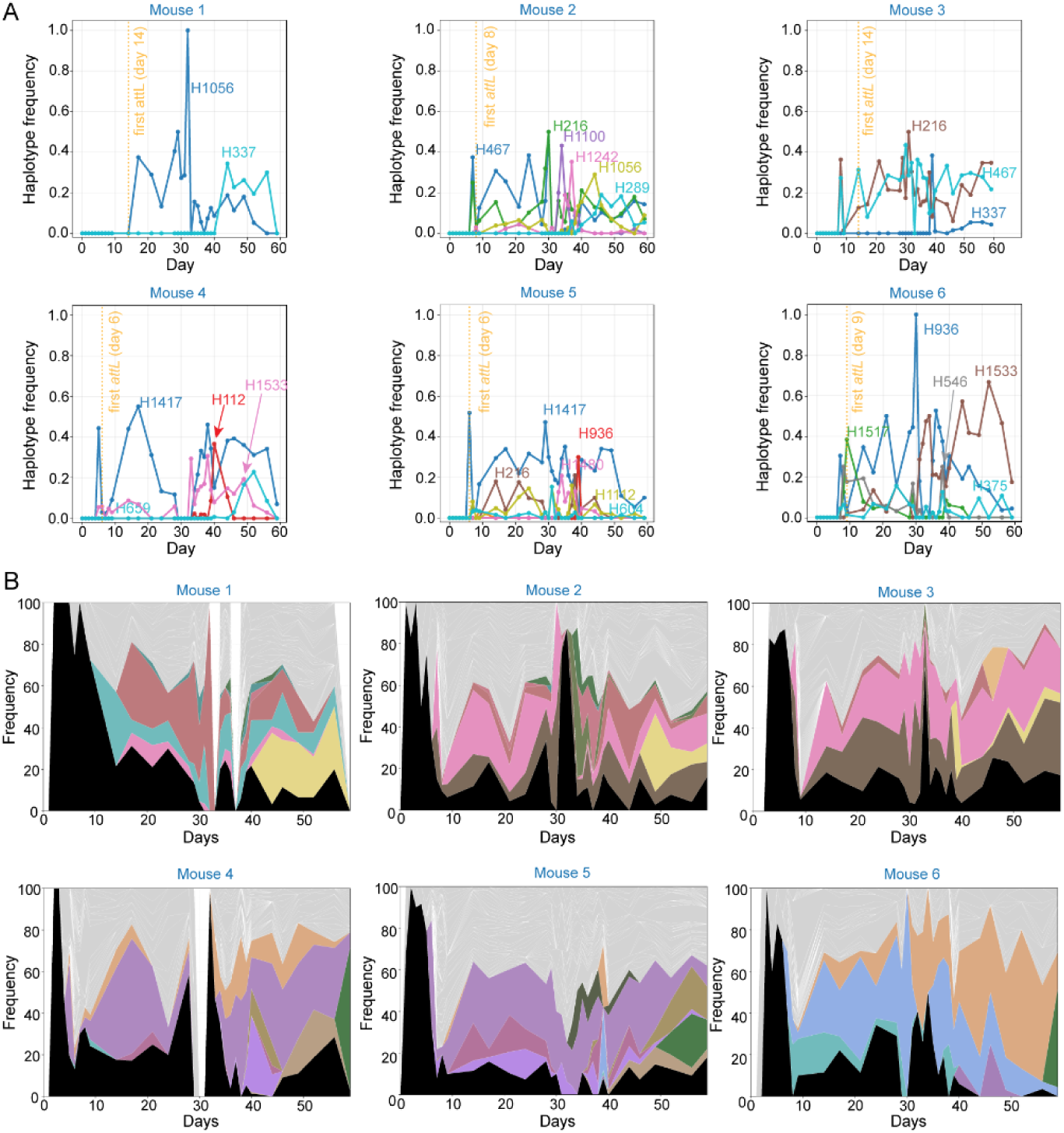
Receptor-binding protein (RBP) haplotypes exhibit reproducible expansion during prophage transfer. A) Longitudinal trajectories of expanding RBP haplotypes in individual SD-fed mice. Expanding haplotypes were defined as non-wild-type haplotypes that increased by ≥15 percentage points from baseline to peak and had a higher mean frequency at the end than at the beginning of the sampling period; wild-type denotes the predominant RBP haplotype at baseline. The orange dotted line indicates the first detection of the recipient-*specific B. intestinalis attL* junction. B) Stacked-area plots showing the frequences of RBP haplotypes over time in individual mice. All haplotypes reaching ≥10% at any time point are shown individually, with rarer variants grouped in gray. The same color denotes the same haplotype across mice; wild-type is shown in black. Mice M1–M3 and M4–M6 were housed together in separate cages. The emergence of shared expanding haplotypes among co-housed mice is consistent with transmission between mice, potentially through coprophagy.

## Materials and Methods

### Defined microbial community assembly

The 38-memember synthetic human gut community was assembled from cultured isolates derived from humanized mice colonized with microbiota from a single human donor (*20*). Briefly, individual strains were grown anaerobically in brain heart infusion medium supplemented with 200 ng/mL tryptophan, 1 mg/mL arginine, 5 ng/mL hemin, 10 ng/mL menadione, 2 mM MgS04, and 100 nM CaCl_2_ (BHI-S) or in a 1:1 mixture of fresh BHI-S and filter-sterilized spent BHI-S from the complete community (BHI-S^sp^). Cultures were cryopreserved in 25% glycerol at –80 °C in plate format. Prior to inoculation, strains were recovered from glycerol stocks in their respective growth medium and grown overnight at 37 °C. Individual strains were then pooled in equal volumes immediately prior to gavage.

### Mouse colonization

Eight- to nine-week-old female germ-free Swiss Webster mice were housed in sterile isolator cages under a 12-h light/dark cycle. All animal procedures were approved by the Stanford University Institutional Animal Care and Use Committee and conducted in accordance with institutional guidelines. Mice were colonized by oral gavage with 200 µL of the pooled community inoculum containing approximately 10^8^ CFU total bacteria. Following colonization, mice were maintained on an autoclaved standard diet (SD; LabDiet 5K67) and co-housed in groups of six for 14 days to allow the community to stabilize.

### Experimental design and dietary intervention

After the 14-day equilibration period, mice were separated into four cages with three mice per cage. Two cages were maintained on SD, while two cages were switched to a microbiota-accessible carbohydrate (MAC)-deficient, high-sugar diet (MD; modified TD.150689, Envigo). The MD contained <1% dietary fiber and was enriched in simple sugars, whereas the SD contained ∼15% complex polysaccharides. Concurrent with the dietary intervention (day 0), all mice were orally gavaged with 200 µL of a barcoded *P. vulgatus* CL09T03C04 transposon insertion library (*35*) containing ∼10^8^ CFU.

Fecal pellets were collected daily during periods of intensive sampling and twice weekly thereafter throughout the 60-day experiment and immediately frozen at –20 °C or –80 °C. On days 28–32, all mice received ciprofloxacin (3 mg per mouse per day) by oral gavage as an antibiotic perturbation. Sampling continued through day 59 post-invasion.

### DNA extraction and metagenomic sequencing

Genomic DNA was extracted from whole fecal pellets using PowerSoil-htp kits (Qiagen) as described previously (*36*). DNA concentrations were quantified in 384-well format using the Quant-iT PicoGreen dsDNA Assay Kit (Thermo Fisher Scientific).

Sequencing libraries were generated in 384-well format using a custom low-volume protocol adapted from the Nextera XT workflow (Illumina), as described previously (*37, 38*). Briefly, genomic DNA samples were normalized to 0.18 ng/µL, and the tagmentation, neutralization, and PCR steps were performed on a Mosquito HTS liquid handler (TTP Labtech). Each reaction contained 0.072 ng of genomic DNA and was amplified for 11 PCR cycles, yielding a final library volume of 4 µL.

Custom 12-bp unique dual indexes were used for sample indexing. The resulting libraries were pooled and purified using AMPure XP beads (Beckman Coulter), targeting a fragment length of approximately 450 bp (∼350-bp insert).

Sequencing was performed on a NovaSeq 6000 using S4 flow cells with 2×150-bp paired-end reads at the Chan Zuckerberg Biohub, targeting 10–20 million paired-end reads per sample. In total, sequencing generated 3.46 billion paired-end reads, corresponding to approximately 1.04 terabases of metagenomic sequence data.

### Metagenomic read quality control and preprocessing

Raw metagenomic sequencing reads were deduplicated using hts_SuperDeduper v1.3.1 (HTStream) to remove PCR duplicates. Adapter sequences were trimmed using Skewer v0.2.2 (*39*) in paired-end mode with the Nextera adapter sequence CTGTCTCTTATACACATCT (-x CTGTCTCTTATACACATCT -m pe). Host-derived reads were removed by mapping the remaining reads to the mouse reference genome (mm39) using Bowtie2 v2.5.4 (*40*) and retaining read pairs for which both mates were unmapped (--un-conc). Host-depleted reads were used for all downstream metagenomic analyses.

### Reference genome sequencing and annotation

High-quality reference genomes were generated for all 38 community members. Individual isolates were cultured anaerobically in BHI-S or 1:1 BHI-S:BHI-S^sp^ according to their strain-specific growth conditions. Genomic DNA was extracted using a modified phenol-chloroform extraction protocol. Long-read sequencing and genomic assembly were performed by Plasmidsaurus (Eugene, OR, USA). Genomes were annotated using Bakta v1.9.1 (*41*). Reference genome sequences are available under NCBI BioProject PRJNA1537695.

### Taxonomic profiling and community composition

Quality-filtered, host-depleted metagenomic reads were taxonomically profiled using using MIDAS v3.0.0 (*42, 43*). Species relative abundances were estimated using the run_species module against a custom database containing reference genomes for the 38 community members.

Samples with fewer than 10 million quality-filtered, host-depleted reads were excluded from community composition analyses. Species relative abundances were extracted from MIDAS output and transformed using the centered log-ratio (CLR) transformation after addition of a pseudocount (10^-6^).

For differential abundance analysis, CLR-transformed abundances were averaged for each mouse within predefined time windows, thereby treating individual mice rather than longitudinal samples as independent observations. Time windows included the post-invasion, pre-antibiotic period (days 7–24), antibiotic treatment (days 28–32), and post-antibiotic recovery (days 33–59). Per-mouse mean CLR abundances were compared between SD-fed and MD-fed mice using Welch’s *t*-test. *P* values were corrected for multiple testing using the Benjamini-Hochberg procedure, and species with an adjusted *q*<0.05 were considered differentially abundant.

### MGE prediction, annotation, and boundary validation

MGE prediction was performed using geNomad v1.5.2 (*44*) to identify plasmids and prophages. Conjugative systems were identified using CONJscan v2.1.1 (*45*) implemented through MacSyFinder, using the CONJscan model set and the ordered_replicon database type. Bakta v1.9.3 annotations were extracted for predicted MGE regions, and MGE annotations were manually curated based on the presence and organization of hallmark genes, including integrases, terminases, capsid proteins, and tail proteins. Protein functions requiring additional validation were analyzed using HHpred (*46*) against the PDB_mmCIF70_20_Feb database.

Because computational MGE predictions do not necessarily resolve precise element boundaries, predicted boundaries were compared with those empirically defined by episomal junctions. For each MGE end, boundary error was defined as the absolute genomic distance between the predicted boundary and the corresponding boundary identified from the episomal junction.

### Detection of circular episomes and empirical MGE boundaries

To identify circular episomal forms generated by excision of iMGEs, we developed a structural-variant (SV) detection approach tailored to metagenomic data. Quality-filtered, host-depleted paired-end reads were mapped to the concatenated 38-genome community reference using Bowtie2 v2.5.4 with default parameters. Alignments were converted to BAM format, sorted, and indexed using SAMtools v1.16.1.

SVs were identified using LUMPY v0.2.13 (*47*) using both discordant paired-end and split-read evidence. Discordant read pairs were extracted using samtools view (excluding reads with flags 1294, i.e., unmapped, mate-unmapped, secondary, duplicate, and concordant paired alignments), and split reads were extracted using the extractSplitReads_BwaMem script distributed with LUMPY. Read groups were added to sorted BAM files using Picard v3.1.1 AddOrReplaceReadGroups, after which BAM files were re-sorted and indexed using SAMtools.

Insert-size distributions were empirically estimated from aligned read pairs using pairend_distro.py, specifying a read length of 146 bp, a maximum distance of 4 standard deviations from the mean (-X 4), and sampling 100,000 read pairs after skipping the first 100,000 alignments to reduce coordinate-order bias. The resulting mean and standard deviation were used to parameterize paired-end evidence for each sample.

LUMPY was run with a minimum weight of 4 (-mw 4) and trim threshold of 0 (-tt 0). Paired-end evidence was supplied with a minimum non-overlap of 101 bp, discordant *z*-score of 5, back distance of 10, weight of 1, and minimum mapping quality of 20. SV calls were output in VCF format.

Circular episomes were identified from duplication (DUP) calls spanning putative iMGE boundaries. Upon excision and circularization, reads spanning the newly formed episomal junction map to opposite boundaries of the chromosomally integrated element, producing a duplication-like SV signature relative to the reference genome. Candidate episomes were restricted to elements 5–500 kb in length. DUP calls were clustered across samples using a 10-kb proximity threshold to identify recurrent calls corresponding to the same putative element. Empirically identified boundaries were compared with computational MGE predictions and manually curated annotations as described above.

### Genome synteny analysis and identification of *de novo* MGE transfer

Pairwise comparisons among the 38 community reference genomes were performed using nucmer from MUMmer v4.0.0 (*48*) with maximal unique match (MUM) clustering. Alignment coordinates were extracted using show-coords with options for reference-based ordering, percent coverage, sequence length, and tab-delimited output (-r -c -l - T).

To distinguish pre-existing MGE-host associations from transfer events occurring during the experiment, we constructed a baseline network from the 38 community reference genomes using the junction-resolved framework described below. This network captured MGE-host associations present in the input community and provided a reference against which integration junctions detected longitudinally *in vivo* were compared. Candidate *de novo* transfer events were defined as MGE-host integration junctions that emerged during the experiment and were absent from both the baseline network and corresponding nucmer alignments.

Candidate events were further filtered to remove technical artifacts and signals arising from homologous sequences. First, events matching baseline junctions were removed using fuzzy coordinate matching; breakend (BND) positions were considered equivalent when they occurred within ±1 kb and involved the same DUP-BND contig pair. Second, nucmer alignments were used to exclude events for which the source MGE and candidate recipient exhibited ≥40% local sequence homology. Local homology was calculated from alignment blocks within a window centered on the BND position and extending 1.25 times the duplication length with an additional 2 kb on each side. Third, recurrent integration signals mapping to the interior of putative MGEs, defined as positions >150 bp from either MGE boundary and observed in ≥2 mice, were excluded as likely arising from homologous MGE sequences rather than independent transfer events. All filtering and network analyses were performed using custom Python scripts available at https://github.com/sariamead/mge_transfer_junctions.

### Junction-resolved quantification of MGE states

For putative MGEs identified from SV analysis, we identified and quantified diagnostic junction sequences corresponding to distinct MGE states. Episomal junctions (*attP*), formed by circularization following excision from the chromosome, were used to quantify the excised MGE state, whereas recipient host-element junctions (*attL*/*attR*) were used to quantify integration into new host genomes. Junction sequences were identified using custom Python scripts based on empirically determined MGE breakpoint coordinates.

For each candidate episome, reads mapping within 1 kb of the predicted breakpoints were extracted from BAM files. Junction-spanning reads were identified from discordant read pairs whose mapping orientations and positions were consistent with circularization. Reads supporting each candidate junction were locally assembled into junction contigs, and junction sequences were validated by remapping metagenomic reads to the assembled junction to confirm read support spanning the breakpoint.

Junction abundance was quantified by counting junction-spanning read pairs and normalizing to the coverage of the corresponding host genome obtained from MIDAS species-level coverage estimates. Junction abundance is therefore reported as junction copies per host genome equivalent. Junction with normalized abundances ≥0.01 copies per host genome equivalent were considered above background, corresponding to approximately one junction-supporting read per bacterial genome equivalent at the sequencing depths analyzed. Temporal dynamics were quantified by tracking normalized junction abundance across longitudinal samples from each mouse.

For MGEs showing evidence of horizontal transfer, recipient-specific integration junctions (*attP*, *attL*/*attR*, and *attB*) were identified and quantified using the same junction-spanning approach. The appearance of recipient-specific *attL*/*attR* junctions that were absent from the baseline community was used as evidence of integration into a new host genome and, after application of the filtering criteria described above, *de novo* horizontal transfer. For mechanistic analyses of the *P. vulgatus* prophage, the episomal *attP* junction and recipient-specific *attL* junction were manually reconstructed from alignment evidence and quantified using the same approach.

All junction extraction, validation, and quantification were performed using custom Python scripts available at https://github.com/sariamead/mge_transfer_junctions.

### MGE single-nucleotide variant analysis

To identify sequence variation within transferring MGEs, quality-filtered metagenomic reads were mapped to MGE reference sequences, and single-nucleotide variants (SNVs) were identified using inStrain v1.9.0 (*49*) with default parameters. SNV allele frequences were tracked longitudinally within individual mice and visualized across MGE coordinates to identify regions exhibiting recurrent sequence diversification.

### Diversity-generating retroelement (DGR) analysis

To characterize diversification within the transferring prophage, we performed longitudinal haplotype analysis of the RBP variable region. Reads spanning the RBP variable region were extracted from BAM files and grouped into haplotypes using custom Python scripts. Because DGR-mediated mutagenesis preferentially introduces substitutions at adenine residues in the template sequence, variants used to define DGR-associated haplotypes were restricted to substitutions corresponding to template A→N changes.

Haplotype frequencies were calculated as the fraction of reads spanning the RBP variable region assigned to each haplotype at each time point. To identify haplotypes whose expansion coincided with prophage integration, we defined an integration window for each mouse extending from 5 days before the first detection of the *B. intestinalis attL* junction to 10 days after the peak *attL* signal. *attL* abundance was quantified using a weighted junction-support score combining directly spanning reads with twice the number of bridge reads. The putative wild-type haplotype, defined as the variant dominant at early time points across mice, was excluded from the analyses.

Haplotypes were classified as transfer-associated (“expanding”) if, within the integration window, their frequency increased by ≥0.15 in absolute frequency, reached a frequency of ≥0.10, and showed a net upward trajectory. To test whether haplotype expansion was temporally associated with prophage integration, we performed a permutation analysis. For each mouse, 10,000 integration windows of matched duration were placed at randomly selected positions within the longitudinal time series, and the number of expanding haplotypes in each window was determined. An empirical *p*-value was calculated by comparing the observed total number of expanding haplotypes across mice with the corresponding null distribution.

Haplotype dynamics were visualized together with species abundance and prophage integration using custom plotting scripts. The relative abundance of each RBP haplotype within the community was estimated by multiplying its within-RBP frequency by *P. vulgatus* relative abundance. Wild-type RBP haplotypes were defined as the most abundant haplotype in early colonization samples independent for each mouse (days ≤3), across all mice.

DGR-associated variable regions and protein domains were predicted using the MyDGR web server (https://omics.informatics.indiana.edu/myDGR/index.php). Protein structures were predicted using AlphaFold2 (*50*) through ColabFold v1.5.2 (*51*). Variant positions were mapped onto predicted structures and visualized using PyMol v2.5 (https://pymol.org).

### Long-read metagenomic sequencing and MGE validation

High-molecular-weight genomic DNA was extracted from fecal samples using a modified protocol combining enzymatic lysis, organic extraction, and selective precipitation. Fecal pellets were resuspended in 200 µL of DNA/RNA Shield (Zymo Research) and incubated overnight at 37 °C with gentle agitation with 100 µg/mL lysozyme (L3790, Millipore Sigma), 200 µg/mL MetaPolyzyme (MAC4LDF-5x1VL, Millipore Sigma), and 200 µg/mL RNase A (19101, Qiagen). Following enzymatic lysis, samples were diluted with 288 µL of EB buffer (10 mM Tris-HCl, pH 8.5) and digested with 400 µg/mL Proteinase K (19131, Qiagen) for 1 h at 56 °C.

DNA was extracted with phenol:chloroform:isoamyl alcohol (25:24:1) with aqueous-phase re-extraction (*52*), followed by extraction with chloroform:isoamyl alcohol (24:1). Contaminating polysaccharides and humic substances were removed using a modified high-salt organic extraction protocol (*53*). Briefly, 398 µL of DNA solution were adjusted to final concentrations of 1 M NaCl and 2 mM EDTA in a total volume of 500 µL and extracted twice with 400 µL of phenol:chloroform:isoamyl alcohol, followed by chloroform:isoamyl alcohol extraction of the pooled aqueous phases.

Polysaccharides were further depleted by fractional ethanol precipitation(*54*). The aqueous phase was mixed with 0.3 volumes of 100% ethanol and centrifuged at 15,000*g* for 15 min at room temperature. The DNA-containing supernatant was transferred to a fresh tube, and the precipitated material was discarded. DNA was subsequently precipitated by adding of 1.7 volumes of 100% ethanol, incubating for 10 min at room temperature, and centrifuging at 16,000*g* for 20 min at 4 °C. DNA pellets were washed twice with 300 µL of 80% ethanol, briefly air-dried, and resuspended in 100 µL of nuclease-free water. Wide-bore pipette tips were used throughout handling of lysates and DNA solutions to minimize shearing. DNA concentration and purity were assessed by Qubit fluorometry and NanoDrop spectrophotometry, respectively. Typical yields were >2 µg, with *A*_260_/*A*_280_>1.8 and *A*_260_/*A*_230_>1.9.

To validate MGE structures and integration events identified by short-read sequencing, fecal samples from SD-fed mice representing time points with high prophage activation or transfer (days 8 and 52) were selected for long-read metagenomic sequencing.

Libraries were prepared using the Oxford Nanopore Technologies Ligation Sequencing Kit (SQK-LSK114-XL) according to the manufacturer’s protocol and sequenced on MinION R10.4.1 flow cells, targeting ≥10 Gb per sample.

Raw POD5 files were basecalled using Dorado v1.3.1 (Oxford Nanopore Technologies) in super-accuracy mode. Basecalled reads were quality-filtered using Filtlong v0.2.1 retaining reads above the 90^th^ percentile (-p 90) for quality. Filtered reads were aligned to a concatenated reference comprising the complete genomes of all 38 members using minimap2 v2.26 (*55*) with parameters -ax map-ont -L. Alignments were converted to BAM format, sorted, and indexed using SAMtools v1.16.1 (*56*).

For the transferring *P. vulgatus* prophage, episomal structures and recipient integration events were validated by identifying long reads spanning the *attP*, *attL*, and *attR* junctions predicted from short-read analysis and confirming the corresponding sequence identity and genomic context. Junction-spanning reads were visualized using custom plotting scripts. For matched samples, short- and long-read junction calls were considered concordant when the same junction was detected by both approaches.

For other MGEs, structural variants (SVs) were identified using Sniffles2 v2.7.5 (*57*) with a minimum SV length of 50 bp (--minsvlen 50), minimum support of three reads (-- minsupport 3), and minimum mapping quality of 20 (--mapq 20). Sniffles was run in reference-aware mode to improve breakpoint resolution. Output VCF files were compressed using bgzip and indexed with tabix for downstream analysis. SVs were clustered and collapsed as described above. Events were retained if they met minimum read-support criteria and, for duplication calls, fell within the target size range of 5–500 kb. Regions flagged as unreliable based on anomalous coverage were excluded. The resulting high-confidence SV set was used for cross-platform concordance analysis.

### Detection of linear extrachromosomal MGEs

In addition to standard SV calling, we developed an approach to identify linear extrachromosomal MGEs from long-read data based on elevated coverage bounded by concentrated read termini, a pattern expected for DNA molecules with defined physical boundaries.

For regions corresponding to duplication calls identified in short-read data, long-read alignments were examined for terminal-read enrichment at the predicted MGE boundaries. Alignment start positions at left boundaries and alignment end positions at right boundaries were quantified within 50-bp windows. Regions with ≥10 terminal reads at both boundaries were classified as candidate linear MGEs.

To identify linear MGEs independently of short-read calls, we also performed a genome-wide scan for coverage-bounded segments. Reference genomes were divided into 500- bp bins, and coverage was smoothed using a three-bin moving average. Candidate segments were initially identified as regions with coverage >1.2-fold above the contig-specific baseline and absolute coverage ≥5X. Candidates were then required to have ≥1.35-fold enrichment relative to 5-kb flanking regions, coverage ≥5X at both boundaries, clusters of read starts at the left boundary and read ends at the right boundary, ≥5X enrichment of termini relative to local background, ≥8 reads at the peak terminal position, and a peak compactness fraction ≥0.25X, defined as the fraction of reads in the boundary cluster occurring at the peak position.

Local background termini density was calculate in 10-kb windows surrounding each boundary, excluding the 1-kb region immediately adjacent to the boundary, using 100-bp bins. This normalization reduced spurious calls from high-coverage regions containing dispersed read termini. Linear MGE calls identified from targeted and genome-wide analyses were merged, and near-duplicate calls with ≥90% reciprocal overlap and boundary differences ≤1 kb were collapsed into single events. When the two approaches produced overlapping calls, targeted calls corresponding to previously identified regions were prioritized.

### Cross-platform concordance

Concordance between short- and long-read SV detection was assessed by comparing genomic coordinates of events identified by the two approaches. Events were considered concordant based on whether they met one of the following criteria: reciprocal coordinate overlap ≥50%, boundary coordinates within 1,000-2,500 bp (parameter dependent), one-sided overlap thresholds (when specified).

### BarSeq processing and fitness analysis

Experiments in this study used a previously described RB-TnSeq mutant library of *P. vulgatus* (*35*). Genomic DNA from fecal and *in vitro* samples containing the barcoded mutant library was processed by barcode sequencing (BarSeq) to quantify the relative abundance of individual transposon mutants. Barcode regions were amplified by PCR using dual-indexed primers to minimize misassignment due to index hopping and sequenced on an Illumina NovaSeq instrument. Sequencing reads were assigned to previously mapped transposon insertions using the barcode-to-gene lookup table generated during library construction. Reads containing unrecognized or ambiguous barcodes were excluded.

Mutant- and gene-level fitness values were calculated as previously described (*23*). Briefly, mutant fitness was calculated as the log_2_ change in barcode abundance in each experimental sample sample relative to the input library. Gene-level fitness was calculated as a weighted average of fitness values from independent transposon insertions assigned to each gene. For the mouse experiments, pre-antibiotic fitness analyses included samples collected through day 28. Statistical significance of gene-level fitness effects was determined as previously described (*23*), with *P* values adjusted for multiple testing using the Benjamini–Hochberg procedure. The same analysis framework was used to quantify fitness across dietary conditions and *in vitro* carbon sources.

### Anaerobic time-lapse microscopy

*P. vulgatus* CL09T03C04 was grown anaerobically in BHI-S supplemented with 1 µg/mL K_3_ at 37 °C. Cells were collected at OD∼1–1.2, washed two times in carbon-free minimal medium, and resuspended in minimal medium containing either glucose or soybean-derived rhamnogalacturonan I (RG-I) as the sole carbon source (5 mg/mL). Cells were transferred to 1.5% agarose pads prepared with the corresponding growth medium and imaged under anaerobic conditions at 37 °C.

Time-lapse phase-contrast images were acquired using a Nikon Ti-E inverted epifluorescence microscope equipped with a 100X (NA 1.4) oil-immersion objective and pco.panda 4.2 aCMOS camera. Images were collected every 10 min for ∼48 h. Propidium iodide (PI; 30 µM) was included in the imaging medium to monitor loss of membrane integrity. Phase-contrast and fluorescence images were acquired at each time point using µManager (*58*).

Cell death was quantified as the fraction of cells becoming PI-positive and/or undergoing lysis during the imaging period. Lysis was defined as [operational definition—e.g., rapid loss of phase-contrast cell structure accompanied by disappearance/collapse of the cell]. The timing of PI uptake and lysis was measured relative to the start of imaging. Three independent biological replicates and twelve fields of view were analyzed for each carbon source.

### Human longitudinal metagenomics analysis

For each donor, a donor-specific reference genome library was constructed from available resolved isolate genomes. Isolate genome assemblies were assessed with CheckM v1.2.3 (*59*) and GTDB-TK v1.0.2 (*60*) to estimate completeness and contamination and to compare reported taxonomic assignments with marker-based lineage assignments. Genomes with inconsistent taxonomic assignments were reclassified where appropriate, while original metadata were retained for traceability.

Representative genomes were selected at the donor–species level. For each species represented among the resolved isolates from a donor, a single reference genome was selected, prioritizing the earliest available isolate followed by assembly quality, including completeness, contamination, genome size, N50, and contig count. For longitudinal human datasets, an early-reference library was constructed separately from species recovered within the earliest available isolate-sampling window for each donor.

Selected representative genomes were concatenated into donor-specific reference FASTA files. Contig headers were renamed to encode source genome and species identity, enabling mapped reads, structural variants, and *nucmer* alignments to be assigned to their isolate genome of origin. A corresponding contig manifest linked each contig to its source genome, curated species assignment, and assembly metadata. Bowtie2 indexes were generated for each concatenated donor-specific reference.

## Data and Code Availability

All code and processed data used to generate figures are available at https://github.com/sariamead/mge_transfer_junctions.

## Acknowledgements

The authors thank the Huang and Sonnenburg labs for helpful discussions. The authors acknowledge support from a Hanna H. Gray Fellowship (to S.M.-M.), NIH RM1 Award GM135102 (to A.D., J.L.S., and K.C.H.), NSF Award EF-2125383 (to K.C.H.), and NIH DP1 DK147449 (to K.C.H.). J.L.S. and K.C.H. are Biohub Investigators. This work was also supported in part by the National Science Foundation under Grant PHYS-1066293 and the hospitality of the Aspen Center for Physics. We thank the Stanford Research Computing Center for use of computational resources on the Sherlock cluster.

